# 17q21.31 sub-haplotypes underlying H1-associated risk for Parkinson’s disease are associated with *LRRC37A/2* expression in astrocytes

**DOI:** 10.1101/860668

**Authors:** KR Bowles, DA Pugh, Y Liu, AE Renton, S Bandres-Ciga, Z Gan-Or, P Heutink, A Siitonen, S Bertelsen, JD Cherry, CM Karch, SJ Frucht, BH Kopell, I Peter, YJ Park, International Parkinson’s Disease Genomics Consortium (IPDGC), A Charney, T Raj, JF Crary, AM Goate

## Abstract

Parkinson’s disease (PD) is genetically associated with the H1 haplotype of the *MAPT* 17q.21.31 locus, although the causal gene and variants underlying this association have not been identified. To better understand the genetic contribution of this region to PD, we fine-mapped the 17q21.31 locus in order to identify novel mechanisms conferring risk for the disease. We identified three novel H1 sub-haplotype blocks across the 17q21.31 locus associated with PD risk. Protective sub-haplotypes were associated with increased *LRRC37A/2* copy number and expression in human brain tissue. We found that LRRC37A/2 is a membrane-associated protein that plays a role in cellular migration, chemotaxis and astroglial inflammation. In human substantia nigra, LRRC37A/2 was primarily expressed in astrocytes, interacted directly with soluble α-synuclein, and co-localized with Lewy bodies in PD brain tissue. These data indicate that a novel candidate gene, *LRRC37A/2*, contributes to the association between the 17q21.31 locus and PD via its interaction with α-synuclein and its effects on astrocytic function and inflammatory response. These data are the first to associate the genetic association at the 17q21.31 locus with PD pathology, and highlight the importance of variation at the 17q21.31 locus in the regulation of multiple genes other than *MAPT* and *KANSL1*, as well as its relevance to non-neuronal cell types.

## INTRODUCTION

The *MAPT* 17q21.31 locus lies within a 1.5Mb inversion region of high linkage disequilibrium (LD), conferring two distinct haplotypes; H1, which has a frequency of ∼0.8 in European ancestry populations, and the less common, inverted H2 haplotype (frequency ∼0.2), which is absent or lower frequency in East and South Asian populations (frequency 0 - 0.09) (Fig 1A). The major haplotype, H1, has been genetically associated with increased risk for multiple neurodegenerative disorders, including *APOE* 4-negative Alzheimer’s disease (AD)^1^, L corticobasal degeneration (CBD)^2^, progressive supranuclear palsy (PSP)^3–5^ and Parkinson’s disease (PD)^6–10^.

**Figure 1.**
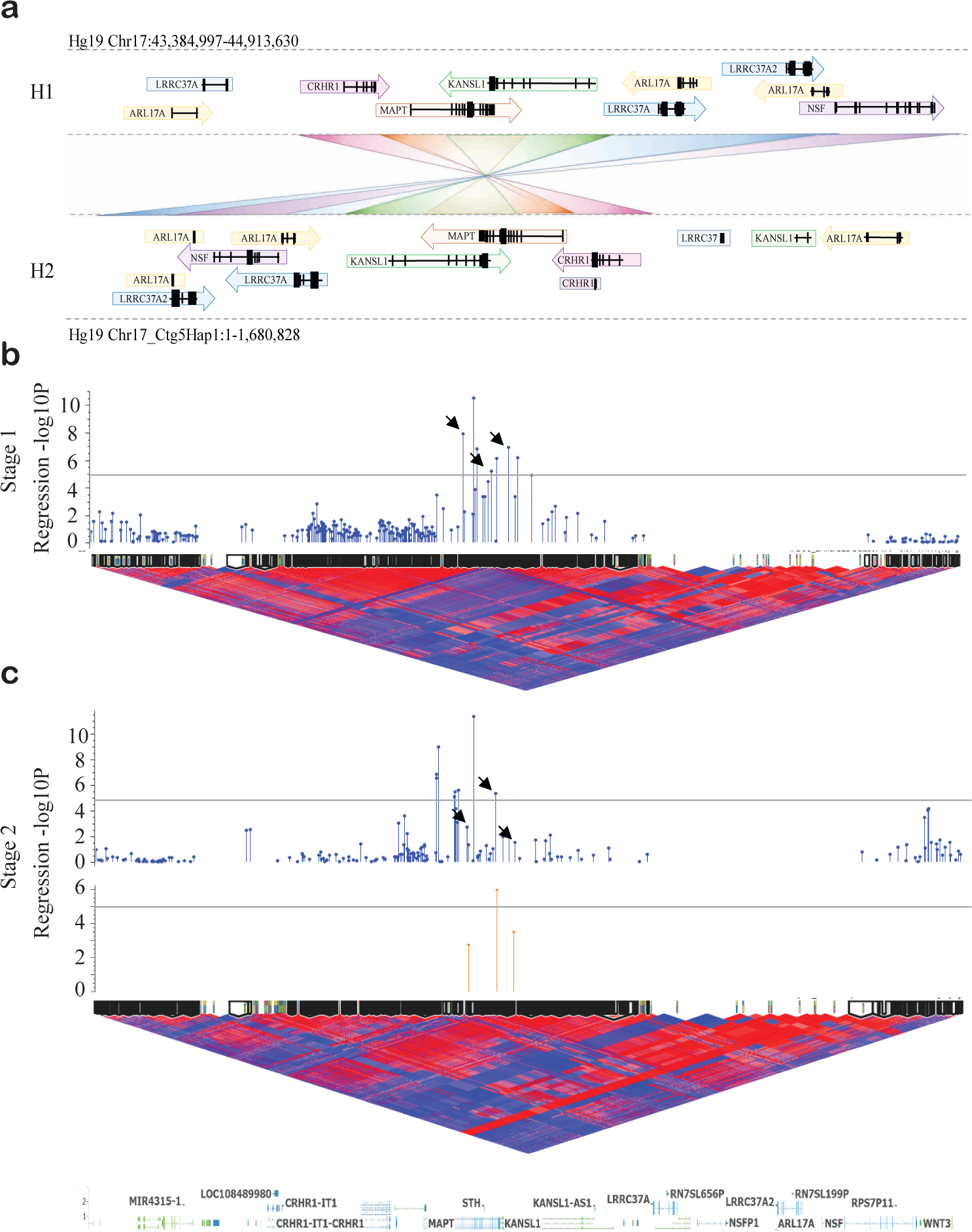
H1 sub-haplotypes within the *MAPT* 17q21.21 inversion region are associated with Parkinson’s disease risk. **A.** Structure of the 17q21.31 locus, which confers two distinct sub-haplotypes defined by gross structural inversion; H1 and H2. Direction of gene orientation in each haplotype is indicated by arrows. Each gene or partial gene is labeled with a distinct color and connected with a crossed rectangle between H1 and H2 to aid visualization of altered gene position between haplotypes. **B-C.** H1 sub-haplotype block association (-log_10_ *p*-value) with PD plotted above H1 homozygote D’ LD structure and sub-haplotype blocks generated from ***B***. Stage 1 data and ***C.*** Stage 2 data, spanning Hg19 Chr17:43384997-44913630. **C**. Association (-log_10_ *p*-value) of blocks calculated in Stage 2 data (blue, top), and H1.1, H1.2 and H1.3 blocks as defined in Stage 1 applied to Stage 2 data (orange, bottom). In LD plots, red indicates high D’ and blue indicates low. Black arrows indicate similar blocks generated across Stage 1 and Stage 2 data. Grey lines indicate genome wide suggestive significance p-value of 1x10^-5^.

PD is a movement disorder that commonly involves executive dysfunction and dementia^11, 12^, but is classically characterized by bradykinesia, tremor, rigidity, postural instability, and numerous non-motor symptoms^13^. Neuropathologically, PD is an - synucleinopathy defined by the presence of intraneuronal accumulation of α-synuclein in Lewy bodies throughout the substantia nigra, brainstem and forebrain^14, 15^. Despite the genetic association with the 17q21.31 *MAPT/*tau locus, aggregated tau is not a typical neuropathological feature of PD^15^, although it is not rare for tauopathy to occur alongside α-synuclein inclusions in α the substantia nigra^16, 17^. There is also no apparent association between PD and the H1c sub- haplotype^18, 19^, which is strongly associated with risk for the primary tauopathies PSP and CBD^2, 3^, indicating that different 17q21.31 locus variants and mechanisms may underlie the relative risk for each disease. Indeed, the 17q21.31 locus spans 1.5Mb including multiple genes in addition to *MAPT*, and comprises several sub-haplotypes, defined by complex rearrangements and copy number variation in their distal regions^20, 21^, the functional impact of which has not yet been explored.

Given the specific loss of midbrain dopaminergic neurons in PD, this cell type has been the main focus of efforts to understand the mechanism of neuronal loss. Similarly, investigation of 17q21.31 variants is also often restricted to neurons, due to the high neuronal expression of *MAPT*. However, a causal role for astrocytes in PD pathogenesis has been recently proposed^22–25^. Indeed, many PD-associated genes are expressed in astrocytes and have known functional roles in astrocyte biology^24^. Furthermore, α-synuclein-positive aggregates have been identified in the α cytoplasm of astrocytes in PD^26–28^, indicating aberrant protein accumulation occurring as a result of either neuronal-glial transmission^22, 28, 29^, or increased expression of endogenous α-synuclein in astrocytes themselves^25^.

Although PD, like PSP and CBD is associated with the H1 haplotype, the odds ratio for risk is substantially lower (1.5 vs 4.5) in PD^2–10^. In contrast to PSP and CBD, PD is not generally associated with the accumulation of tau. Together these observations suggest that the PD genetic association with the 17q21.31 locus may be driven by genes other than *MAPT*. Here, we report three sub-haplotype blocks associated with PD risk within the 17q21.31 H1 haplotype clade, with both protective and risk-associated sub-haplotypes within each block. We show that protective sub-haplotypes are associated with increased expression and copy number of *LRRC37A/2*, which we demonstrate is an astrocyte-enriched membrane-associated protein with a role in chemotaxis and the inflammatory response, and co-localizes with both soluble –α synuclein and Lewy bodies in human substantia nigra. These findings link the genetic association at the 17q21.31 locus with PD pathology, and support the hypothesis of astroglial dysfunction as a key contributing factor to PD disease pathogenesis.

## RESULTS

### PD risk is associated with the 17q21.31 H1 haplotype

To confirm the genetic association of the 17q21.31 H1 haplotype with PD risk, we carried out a case-control association analysis across the region of interest (Figure S1A-E, Table S1) in two independent datasets (Stage 1 2,780 PD cases, 6,384 controls; Stage 2 2,699 cases, 2,230 controls; Table S2). The SNP with the strongest association in Stage 1 (rs17763050, *p* = 2.74 x10^-9^) was in high LD with the known H1/H2 haplotype tag SNP rs8070723 (D’ = 0.98, Figure S1B-C, F). Both SNPs were associated with odds ratios (ORs) ∼0.8 (95% CI ± 0.1; Figure S1B-C, Table S1), consistent with previously reported effect sizes^6, 8^. Due to a smaller cohort size and consequent lack of power, the association with 17q21.31 was less prominent in Stage 2 data (Figure S1D-E, Table S1), and the top SNP did not tag the H2 haplotype. However, meta-analysis of both cohorts confirmed a significant association between PD risk and the 17q21.31 locus (Figure S1B-C, Table S1), with the H2 haplotype conferring protection (OR 0.82 (95% CI 0.76-0.89)) and the H1 haplotype therefore associated with increased risk.

As the major H1 haplotype was associated with increased risk for PD, we repeated the association analysis in H1 homozygotes alone in order to identify variants of H1 that may confer additional risk for PD (Figure S1G-J, Table S1). While association across the 17q21.31 locus was weaker in H1 homozygotes compared to the full data set, we observed a distinct signal spanning *MAPT* and *KANSL1* in Stage 1 and Stage 2 analyses (Figure S1G-H). The Stage 1 top SNP, rs41543512, was associated with an OR of 1.21 (95% CI 1.10-1.32, *p*<0.001; Figure S1I, Table S1), but did not reach statistical significance in Stage 2 data or meta-analysis (Table S1). The most significant SNP in the Stage 2 analysis (rs139217062) was not present in Stage 1 data, although the second most significant variant from Stage 2, rs16940711, was not significant in Stage 1 data or by meta-analysis (Table S1, Figure S1J). These data suggest that the H1 association with PD may be more complex than variation in individual SNPs.

### 17q21.31 H1 sub-haplotype blocks spanning *MAPT* & *KANSL1* are associated with PD risk

As SNP-based association analyses were unsuccessful in identifying significant H1 sub- haplotypic variants contributing to PD risk, we decided to leverage the presence of high linkage disequilibrium (LD) within this region to investigate sub-haplotype blocks. We calculated discrete sub-haplotype blocks spanning the 17q21.31 locus using the D’ measure of LD, and performed a logistic regression association analysis on each block (Figure 1B-C, Table 1). This approach greatly improved the power to detect disease-associated H1 variants in both Stage 1 and Stage 2 data. In Stage 1, we observed a peak spanning *MAPT* and the first 5 exons of *KANSL1* (Figure 1B) that reached the genome-wide suggestive significance threshold of *p*=1x10^- 5^. Within this peak, three sub-haplotype blocks showed substantial overlap of SNPs in independently calculated blocks in Stage 2 data (Stage 1 blocks H1.1 (*p* = 1.73x10^-6^), H1.2 (*p* = 2.4x10^-4^) and H1.3 (*p* = 1.05x10^-5^); Figure 1C, Table 1). We then applied the Stage 1-constructed blocks to Stage 2 data and observed replication of their association with PD risk (Figure 1C, Table 1). Block H1.2 was highly significant in Stage 2 data (*p* = 1.12x10^-9^), while both blocks H1.1 and H1.3 were nominally significant (*p* <0.002 and *p* <0.0003, respectively).

**Table 1.**
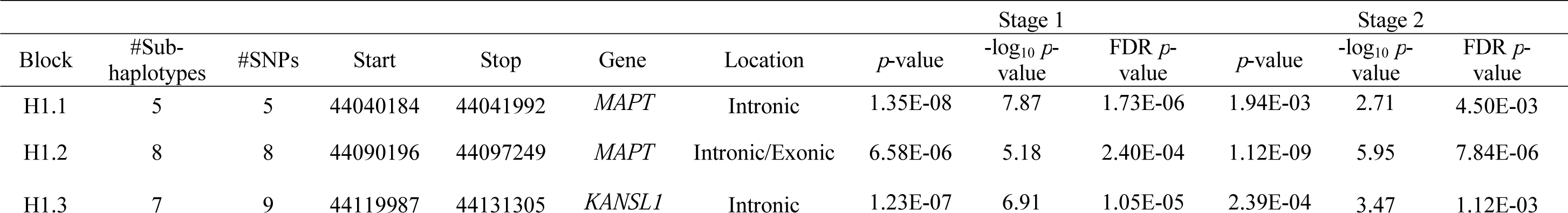
17q21.31 H1 sub-haplotype blocks associated with PD susceptibility.

Blocks H1.1, H1.2 and H1.3 each consist of multiple SNPs in high LD (Figure S1K) that generate multiple sub-haplotypes (Figure S2A-F). Each sub-haplotype was differentially associated with PD susceptibility (Figure S2A,C,E, Table 2), with each block containing both risk- and protective sub-haplotypes with ORs ranging from 0.37 (95% CI 0.32-0.42, *p*=1.3x10^-49^, H1.1c) to 2.51 (95% CI 2.2-2.86, *p*=2.4x10^-45^, H1.1e). Despite the presence of significant heterogeneity between Stages 1 and 2, the most frequently occurring sub-haplotypes were replicated in both analysis stages and by fixed effects meta-analysis (Table 2). Specifically, we identified two sub-haplotypes in block H1.1; H1.1b and H1.1e, which increased risk in both datasets when compared against the most common haplotype, with ORs ranging from 1.26-1.6 (fixed effects *p*<0.001) and 1.45-2.51 (fixed effects *p*<0.001), respectively (Figure S2A-B, Table 2). In the same block we also observed a protective sub-haplotype (H1.1c), associated with an OR ranging 0.37-0.96 (fixed effects *p*<0.001; Figure S2A-B, Table 2). Blocks H1.2 and H1.3 encompassed multiple sub-haplotypes with frequencies < 0.1, and exhibited greater variability and heterogeneity across stages. However, we identified one risk-associated sub-haplotype in block H1.2 (H1.2c, OR = 1.12-1.31, fixed effects *p*<0.01) and two protective sub-haplotypes in block H1.3 (H1.3b; OR = 0.95-0.43, fixed effects *p*<0.001, H1.3g; OR = 0.55-0.83, fixed effects *p*<0.002; Table 2, Figure S2C-F). These data reflect the variability and complexity present across the 17q21.31 locus, and indicate that numerous H1 variants exist that are associated with differential levels of risk for PD.

**Table 2.** H1 sub-haplotypes associated with PD susceptibility.

| Block | Sub-haplotype ID | Sub-haplotype | Stage 1 |  |  | Stage 2 |  |  | Meta |  |  |  |  |  |
| --- | --- | --- | --- | --- | --- | --- | --- | --- | --- | --- | --- | --- | --- | --- |
| | | | Frequency (Case/Control) | OR (95% CI) | OR Fisher's exact $p^{sig}$ | Frequency (Case/Control) | OR (95% CI) | OR Fisher's exact $p^{sig}$ | RE OR (95% CI) | RE Heterogeneity ($p$) | RE $p^{sig}$ | FE OR (95% CI) | FE Heterogeneity ($p$) | FE $p^{sig}$ |
| H1.1 | H1.1a | ACTCT | 0.25 (0.26/0.24) | 1 | - | 0.21 (0.2/0.23) | 1 | - | 1 | - | - | 1 | - | - |
|  | H1.1b | ACTTG | 0.27 (0.38/0.22) | 1.6 (1.45-1.78) | 2.04E-19*** | 0.38 (0.39/0.36) | 1.26 (1.09-1.46) | 2.0E-03** | 1.43 (1.13-1.81) | 7.06 (0.0079) | 3.0E-03** | 1.48 (1.36-1.61) | 7.06 (0.0079) | 0*** |
|  | H1.1c | GCCTG | 0.18 (0.09/0.23) | 0.37 (0.32-0.42) | 1.3E-49*** | 0.14 (0.13/0.16) | 0.96 (0.79-1.15) | 6.40E-01 | 0.59 (0.23-1.52) | 67.79 (<0.0001) | 2.70E-01 | 0.51 (0.46-0.57) | 67.85 (<0.0001) | 0*** |
|  | H1.1d | ATCTG | 0.18 (0.06/0.24) | 0.22 (0.19-0.26) | 6.5E-92*** | 0.23 (0.24/0.21) | 1.29 (1.1-1.52) | 2.0E-03** | 0.54 (0.1-3.01) | 231.51 (<0.0001) | 4.90E-01 | 0.49 (0.44-0.55) | 232.74 (<0.0001) | 0*** |
|  | H1.1e | ACCTG | 0.12 (0.21/0.08) | 2.51 (2.2-2.86) | 2.4E-45*** | 0.02 (0.02/0.02) | 1.45 (0.98-2.17) | 5.70E-02 | 1.97 (1.15-3.36) | 7.13 (0.0079) | 1.3E-02* | 2.37 (2.1-2.68) | 7.13 (0.0076) | 0*** |
| H1.2 | H1.2a | TTTCGATG | 0.48 (0.44/0.5) | 1 | - | 0.49 (0.49/0.48) | 1 | - | 1 | - | - | 1 | - | - |
|  | H1.2b | TCTCGATG | 0.17 (0.18/0.16) | 1.27 (1.14-1.4) | 2.02E-05*** | 0.17 (0.17/0.18) | 0.93 (0.8-1.09) | 3.56E-01 | 1.09 (0.81-1.47) | 10.57 (0.001) | 5.70E-01 | 1.14 (1.04-1.24) | 10.58 (0.0011) | 3.0E-03** |
|  | H1.2c | TTAAAATA | 0.15 (0.16/0.14) | 1.31 (1.16-1.47) | 4.52E-06*** | 0.19 (0.21/0.18) | 1.12 (0.97-1.3) | 1.21E-01 | 1.22 (1.05-1.42) | 2.65 (0.103) | 9.0E-03** | 1.23 (1.13-1.35) | 2.65 (0.1037) | 4.41E-06*** |
|  | H1.2d | TTAAGATG | 0.07 (0.08/0.06) | 1.36 (1.16-1.59) | 1.2E-04*** | 0.04 (0.03/0.05) | 0.52 (0.38-0.69) | 3.46E-06*** | 0.84 (0.33-2.19) | 35.25 (<0.0001) | 7.27E-01 | 1.07 (0.94-1.23) | 35.28 (<0.0001) | 2.90E-01 |
|  | H1.2e | CTTCGATG | 0.06 (0.06/0.06) | 1.17 (0.99-1.39) | 5.64E-02 | 0.08 (0.08/0.08) | 1.05 (0.85-1.3) | 6.76E-01 | 1.12 (0.99-1.28) | 0.72 (0.40) | 7.00E-02 | 1.12 (0.99-1.28) | 0.72 (0.397) | 7.00E-02 |
|  | H1.2f | TTTCGGTG | 0.05 (0.05/0.05) | 1.16 (0.96-1.39) | 1.18E-01 | 0.02 (0.02/0.03) | 0.58 (0.41-0.83) | 7.1E-03** | 0.83 (0.42-1.64) | 12.41 (0.0004) | 6.00E-01 | 0.99 (0.84-1.16) | 12.41 (0.0004) | 9.10E-01 |
|  | H1.2g | TTAAAATG | 0.01 (0.01/0.01) | 1.1 (0.77-1.54) | 6.07E-01 | - | - | - | - | - | - | - | - | - |
|  | H1.2h | TTTCGACG | 0.01 (0.01/0.01) | 1.28 (0.89-1.84) | 1.64E-01 | - | - | - | - | - | - | - | - | - |
| H1.3 | H1.3a | GACTGAGAT | 0.28 (0.32/0.27) | 1 | - | 0.32 (0.32/0.32) | 1 | - | 1 | - | - | 1 | - | - |
|  | H1.3b | CATTAGGGC | 0.18 (0.11/0.22) | 0.43 (0.38-0.49) | 1.66E-41*** | 0.18 (0.18/0.19) | 0.95 (0.81-1.11) | 5.22E-01 | 0.64 (0.29-1.44) | 58.53 (<0.0001) | 2.80E-01 | 0.57 (0.52-0.63) | 58.57 (<0.0001) | 0*** |
|  | H1.3c | GATTGAGAT | 0.17 (0.21/0.16) | 1.11 (0.99-1.24) | 7.37E-02 | 0.16 (0.16/0.17) | 0.95 (0.80-1.12) | 5.33E-01 | 1.04 (0.89-1.21) | 2.24 (0.134) | 6.10E-01 | 1.06 (0.96-1.16) | 2.24 (0.1343) | 0.23 |
|  | H1.3d | CTTTGGTGC | 0.15 (0.12/0.16) | 0.61 (0.54-0.7) | 1.58E-14*** | 0.2 (0.21/0.18) | 1.16 (0.99-1.36) | 5.81E-02 | 0.84 (0.45-1.56) | 35.27 (<0.0001) | 5.80E-01 | 0.77 (0.7-0.85) | 35.28 (<0.0001) | 2.96E-07*** |
|  | H1.3e | CATTGGGGC | 0.09 (0.14/0.07) | 1.67 (1.45-1.92) | 2.79E-13*** | 0.02 (0.02/0.02) | 0.81 (0.54-1.21) | 2.83E-01 | 1.19 (0.58-2.42) | 12.01 (0.0004) | 6.30E-01 | 1.54 (1.35-1.75) | 12.01 (0.0005) | 4.57E-11*** |
|  | H1.3f | CATTGGGGT | 0.08 (0.07/0.09) | 0.6 (0.51-0.70) | 9.34E-11*** | 0.09 (0.1/0.08) | 1.15 (0.93-1.42) | 2.00E-01 | 0.81 (0.44-1.5) | 21.37 (<0.0001) | 5.10E-01 | 0.74 (0.66-0.84) | 21.38 (<0.0001) | 3.59E-06*** |
|  | H1.3g | GATCGAGAT | 0.04 (0.04/0.04) | 0.83 (0.67-1.03) | 9.39E-02 | 0.03 (0.02/0.02) | 0.55 (0.38-0.78) | 5.58E-04*** | 0.7 (0.49-1.02) | 3.48 (0.062) | 6.60E-02 | 0.75 (0.63-0.9) | 3.48 (0.062) | 1.6E-03** |

**OR** = Odds ratio; **CI** = Confidence interval; **RE** = Random effects meta-analysis; **FE** = Fixed effects meta-analysis. \* $p<0.05$ , \*\* $p<0.01$ , \*\*\* $p<0.001$

### PD-associated sub-haplotypes are associated with *LRRC37A/2* gene expression in human brain

In order to elucidate the functional consequences of PD-associated H1 block sub-haplotypes we queried publicly available post-mortem human brain RNA-seq data from dorsolateral prefrontal cortex (PFC) and temporal cortex (TCX) from the AMP-AD and CommonMind consortia (Table S3). Despite the position of the H1 association peaks across *MAPT* and *KANSL1*, we did not observe any differences in the expression of either of these genes between any sub-haplotypes in any block (Figure 2C, Figure S3A-C). The only genes within the 17q21.31 locus that had a significant association with H1 PD-associated sub-haplotypes were *LRRC37A* and its paralog *LRRC37A2* (a.k.a *LRRC37A/2*; Figure 2A-B, Figure S3D). We observed significantly increased *LRRC37A*/*2* expression in protective sub-haplotypes, specifically in H1.1c and H1.3b (H1.1c ∼ 4.7 fold, *p*<0.001, H1.3b ∼ 5 fold, *p*<0.001), as well as in sub-haplotypes whose effects were not replicated across PD data-sets; H1.2b and H1.3e (H1.2b ∼5, *p*<0.001, H1.3e ∼2.9x, *p*<0.01), but were protective in the Stage 2 PD analysis (Figure 2A-B). qRT-PCR on postmortem prefrontal cortex from a small number of individuals supported the observation of increased *LRRC37A/2* expression in these sub-haplotypes (Figure S3D).

**Figure 2.**
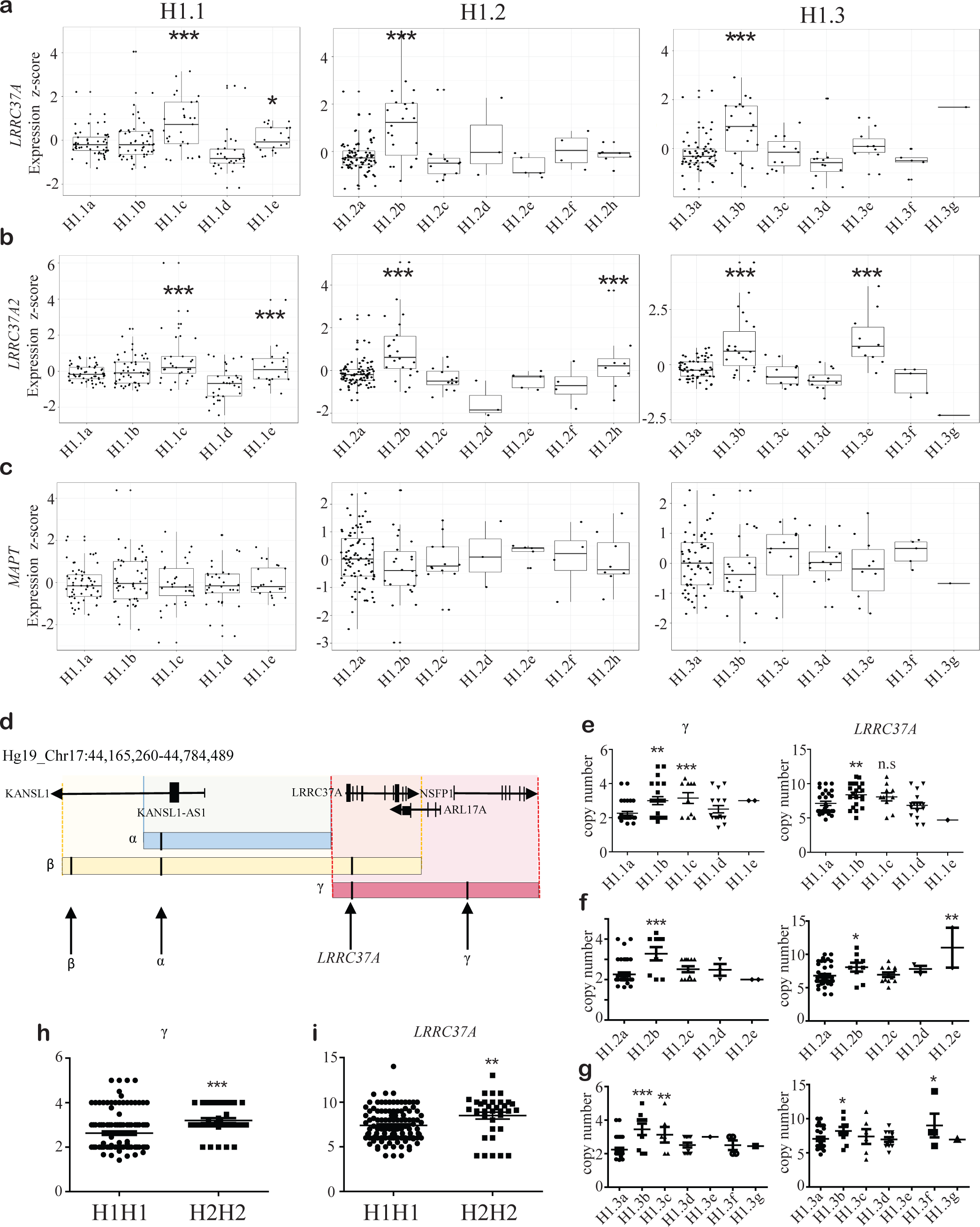
PD-associated sub-haplotypes are associated with *LRRC37A/2* expression and copy number. **A-C**. Expression of ***A***. *LRRC37A, **B.** LRRC37A2* and ***C.*** *MAPT* in human brain tissue, measured by RNA-seq across three different cohorts, split by sub-haplotype in blocks H1.1, H1.2 and H1.3. **D.** Schematic of the regions of copy number variation in the 3’ distal end of the 17q21.31 locus, as defined by Boettger et al 2012^20^. Black arrows indicate the location of dPCR probes for (left to right) beta, alpha, *LRRC37A* and gamma. **E-G**. Copy number of gamma and *LRRC37A/2* regions in blocks ***E.*** H1.1, ***F.*** H1.2 and ***G.*** H1.3. **H-I**. Copy number of ***H.*** gamma and ***I.*** *LRRC37A/2* regions between H1 and H2 homozygotes. All statistical comparisons are against the most common sub-haplotype. ns = not significant, \**p* < 0.05, \*\**p* < 0.01, \*\*\**p* < 0.001.

We observed a significant reduction in *LRRC37A/2* expression in the PD risk-associated H1.1b sub-haplotype by qRT-PCR in human brain tissue (*p*<0.01; Figure S3D), but not in the RNA-seq data. We also did not observe any difference in *MAPT* exons 2, 3 or 10 percent spliced in (PSI) values between sub-haplotypes (Figure S4A). Interestingly, *LRRC37A/2* expression was significantly higher in the protective 17q21.31 H2 haplotype (Figure S4B), which is consistent with our observation of increased *LRRC37A/2* expression in protective H1 sub-haplotypes. These data suggest that both the 17q21.31 H1/H2 and H1 sub-haplotype genetic associations with PD risk may be due to variable *LRRC37A/2* expression.

### Protective sub-haplotypes are associated with increased *LRRC37A/2* copy number

The 17q21.31 locus is structurally complex and encompasses regions of copy number variation (CNV) at its distal ends^20, 21^. As sub-haplotype blocks within *MAPT* and *KANSL1* were associated with altered *LRRC37A/2* expression, we tested whether they also tagged structural variants^20^ in individuals homozygous for sub-haplotypes of interest (Figure 2D-H, Figure S5A- D). Using DNA derived from either blood or brain tissue (Table S3) we performed digital PCR (dPCR) for *MAPT*, alpha, beta and gamma regions (Figure 2D)^20^, as well as for *LRRC37A/2* specifically (Figure 2E-G, Figure S5A-D).

The majority of the structural variation in alpha and beta regions was found in the most common sub-haplotype for each block (H1.1a, H1.2a and H1.3a), with each subsequent sub-haplotype carrying fewer copies of these regions (Figure S5A-C). However, gamma and *LRRC37A/2* CNVs varied by sub-haplotype within each block; those sub-haplotypes exhibiting increased *LRRC37A/2* expression were also associated with significantly increased gamma (H1.1c R^2^=0.23, H1.2b R^2^=0.25, H1.3b R^2^=0.34, ∼3-5 copies compared to 2-3 copies in controls, *p*<0.05) and/or *LRRC37A/2* copy number (H1.1c R^2^=0.06, H1.2b R^2^=0.08, H1.3b R^2^=0.09, ∼ 8-11 copies compared to 5-10 copies in controls, *p*<0.05; Figure 2E-G). Interestingly, risk-associated sub-haplotype H1.1b was associated with increased gamma and *LRRC37A/2* copy number (*p*<0.01; Figure 2E) but not *LRRC37A/2* expression (Figure 2A-B), indicating that additional factors likely contribute to *LRRC37A/2* expression and PD risk in this locus, such as chromatin looping or variants within *LRRC37A/2* itself. Consistent with the expression data, we also observed increased beta, gamma and *LRRC37A/2* copy number in H2 homozygotes compared to H1 (Figure S5D). These data suggest that structural variation at the distal end of the 17q21.31 locus may underlie increased expression of *LRRC37A/2* in protective H2 and H1 sub-haplotypes.

### *LRRC37A* is a membrane-associated protein implicated in cellular migration, chemotaxis and inflammation

As PD sub-haplotypes converge on the expression and/or copy number of *LRRC37A/2*, we explored the likely function of this gene. We carried out RNA-seq analysis in HEK293T cells overexpressing *LRRC37A/2* in order to mimic increased copy number and to elucidate a potential function for this gene. The number of significantly differentially expressed protein-coding genes (fold change ± 1.5, adjusted *p*<0.05) in the context of *LRRC37A/2* overexpression was minimal (28 upregulated, 21 downregulated), suggesting that *LRRC37A/2* is unlikely to play a major regulatory role. In order to confirm that we were not observing spurious changes in gene expression due to gross overexpression in a cell culture model, we carried out a titration of *LRRC37A/2* overexpression in HEK293T cells. We observed dose-dependent changes in the expression of genes that were significantly up or downregulated in our RNA-seq data (Figure S6B-F), confirming that the expression of these genes is likely to be altered by *LRRC37A/2* expression.

Functional enrichment of gene ontology (GO) terms for significantly differentially expressed genes indicated a role for LRRC37A/2 at the cell membrane (Figure 3A-B), which we confirmed by western blot in HEK293T cells (Figure S6A). In addition, *LRRC37A/2* overexpression also resulted in significant enrichment for *cell communication* (GO:0007154, *p*<0.05) and *neuroactive ligand-receptor interaction* (KEGG:04080, *p*<0.05) pathways, as well as nominal enrichment for membrane-component-related pathways (Figure 3B).

**Figure 3.**
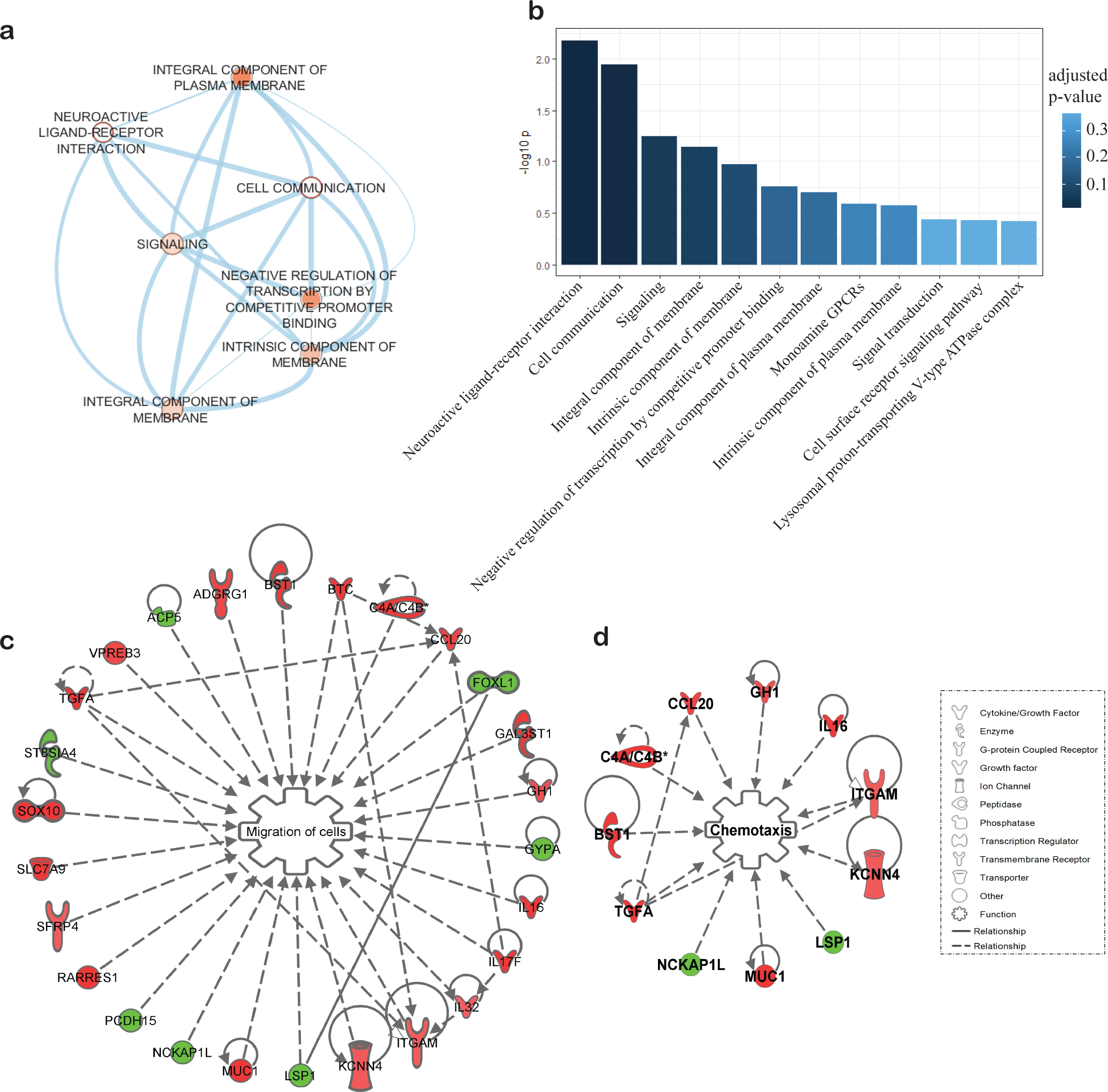
*LRRC37A/2* expression is associated with increased cellular migration, chemotaxis and inflammation. **A.** Enriched GO terms for significantly differentially expressed genes following *LRRC37A/2* overexpression in HEK293T cells. Paler node colors indicate less significant enrichment *p*- values, and edge thickness indicates the proportion of shared genes between GO terms. **B.** -log_10_ adjusted *p*-values for top 12 functionally enriched gene ontology terms for significantly differentially expressed genes in *LRRC37A*/2-overexpressing cells. **C-D.** Significantly enriched ***C.*** *Migration of cells* and ***D.*** *Chemotaxis* pathways associated with *LRRC37A/2* overexpression, derived from Ingenuity pathway analysis. Red genes indicate upregulation, green indicates downregulation

Ingenuity pathway analysis (IPA) also suggested a role for LRRC37A/2 at the plasma membrane and in the extracellular space; specifically, these analyses indicated that increased *LRRC37A/2* expression resulted in upregulated cellular movement pathways, such as increased *migration of cells* (*p*<0.01, z score = 1.85; Figure 3C) and upregulation of *chemotaxis* (*p*<0.05, z score = 0.918; Figure 3D). Several upregulated genes within these pathways are also essential in regulation of the inflammatory response; both *IL17F* and *IL32* are pro-inflammatory cytokines that mediate the inflammatory response of astrocytes^30, 31^, whereas *IL16* acts as a chemoattractant for cells expressing CD4. These data therefore suggest that increased *LRRC37A/2* expression may mediate astroglial inflammation and modify cellular migration in response to a stimulant, such as α-synuclein.

### LRRC37A/2 is expressed in astrocytes and interacts with soluble and aggregated synuclein

Our pathway analysis data indicated that *LRRC37A/2* was associated with cellular signaling and pro-inflammatory pathways relevant to astroglial function. We therefore sought to confirm whether *LRRC37A/2* was expressed in these cells. As *LRRC37A/2* expression and copy number was higher in 17q21.31 H2 haplotype carriers compared to H1, and due to difficulty in identifying multiple human iPSC lines with specific H1 sub-haplotypes, we compared the expression of *LRRC37A/2* and genes altered by *LRRC37A/2* expression in iPSC-derived neurons and astrocytes homozygous for either H1 or H2 haplotypes in order to identify the most relevant neural cell type (Figure 4A, Figure S6G). While *LRRC37A/2* was expressed in both cell types, we observed increased expression of *LRRC37A/2* and associated genes in H2 astrocytes, but not in H2 neuronal cultures (Figure 4A, Figure S6G), suggesting that H1/H2-associated *LRRC37A/2* expression changes may be specifically impacting astroglial gene expression and function. We also confirmed that LRRC37A/2 was present in the plasma membrane in both iPSC-derived neurons and astrocytes by isolating cytosolic and membrane-associated proteins from each cell type and analyzing the resulting fractions by western blot (Figure 4B-C).

**Figure 4.**
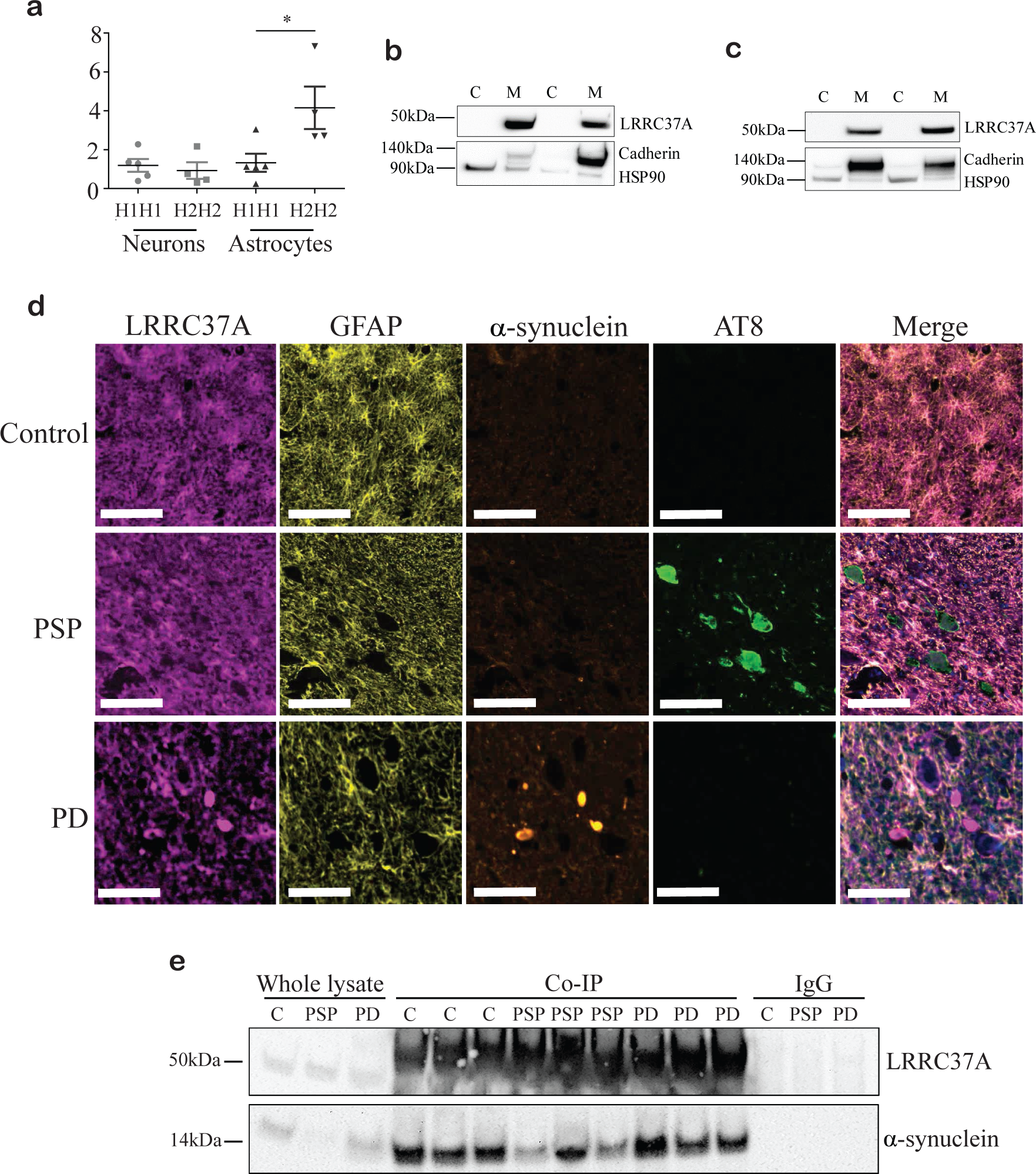
LRRC37A/2 is a membrane-associated protein expressed in astrocytes and co-localized with α-synuclein. **A.** qRT-PCR for *LRRC37A* expression in H1 and H2 homozygote iPSC-derived neurons and astrocytes. N=3 in duplicate. ns = not significant, \**p* < 0.05. **B-C.** Western blots for LRRC37A/2 in cytosolic (C) and membrane (M) fractions from ***B.*** iPSC- neurons and ***C.*** iPSC-astrocytes. Cytosolic fractions were confirmed by labelling with an anti- HSP90 antibody, and membrane fractions were confirmed by labelling with an anti-Pan- Cadherin antibody, N=6. **D.** Representative images from multiplex immunofluorescent staining of human control, PSP and PD substantia nigra sections with astrocyte marker GFAP, LRRC37A, α-synuclein and pathologically phosphorylated Tau (AT8). Scale bar = 100µm, N = 4-5. **E**. Co-immunoprecipitation (Co-IP) of LRRC37A (top panel) with soluble α-synuclein (bottom panel) in substantia nigra from control (C), PSP and PD brain, N=3. Whole protein lysates and IgG only controls were included for comparison.

In order to assess whether LRRC37A/2 was expressed in mature astrocytes in human brain tissue, we carried out multiplex immunofluorescence staining in human substantia nigra from PD, PSP and aged controls (Figure 4D). We found that in all cases, LRRC37A/2 co- localized with the astrocyte marker GFAP, but not the microglia marker IBA1 (Figure 4D, Figure S6H). In contrast, α-synuclein positivity was observed only in PD substantia nigra, and α hyperphosphorylated tau (labeled with AT8) was present only in PSP brain (Figure 4D). Interestingly, in regions with Lewy body pathology there was reduced staining intensity of LRRC37A/2 in astrocytes, and colocalization of LRRC37A/2 with Lewy bodies Figure 4D). ( However, there was no association between tau AT8 positivity and LRRC37A/2 expression in PSP substantia nigra (Figure 4D), indicating that LRR37A/2 accumulation is specific to PD pathology. To validate the association of LRRC37A/2 with α-synuclein, we performed co-α immunoprecipitation from control, PSP and PD substantia nigra tissue (Figure 4E). We found that soluble α-synuclein bound LRRC37A/2 in all cases, whereas IgG alone did not (Figure 4E), α indicating that LRRC37A/2 and α-synuclein likely form a normal complex in human brain that α becomes disordered in the context of PD pathogenesis. These data are not only the first to identify a role for LRRC37A/2 in astrocytes, but are the first to link the genetic association at the 17q21.31 locus with PD pathology.

## DISCUSSION

By constructing discrete sub-haplotype blocks across the 17q21.31 H1 locus, we have identified multiple novel H1 sub-haplotypes associated with variable levels of PD risk. The *MAPT* gene, encoding the microtubule associated protein tau, is central to this locus and is likely the causal gene for other neurodegenerative disorders genetically associated with the H1 haplotype, such as PSP and CBD^2, 3^, given that they are characterized neuropathologically by tau hyperphosphorylation and accumulation. However, while tau pathology can occur alongside –α synuclein inclusions in the substantia nigra^16, 17^, it is not a typical neuropathological feature of PD^15^. The causal gene underlying the genetic association across the 17q21.31 locus with PD risk has therefore been unclear, although a recent study proposed variants in *KANSL1* as underlying this association by altering mitophagy^32^. In contrast, we did not find any effect of our PD-associated H1 sub-haplotypes on either *KANSL1* expression or copy number, but did observe a consistent association with the expression of another gene in the locus, *LRRC37A/2*. However, these data do not rule out that *KANSL1* variants may alter disease risk independently of *LRRC37A/2* expression between the major haplotype clades and H1 sub-haplotypes.

We found that protective H1 sub-haplotypes were associated with increased expression of *LRRC37A/2*. This is consistent with increased expression of these genes in the protective H2 haplotype, suggesting that there is likely a shared mechanism of protection from PD between H2 and specific sub-haplotypes of H1. Furthermore, analysis of CNVs in the 3’ distal end of the 17q21.31 locus suggested that protective sub-haplotypes were tagging structural variants defined by increased gamma region^20^ and *LRRC37A/2* copy number, which likely underlies the increased expression of these genes. *LRRC37A* is a core duplicon on chromosome 17^33^ and is present at the inversion breakpoint of the 17q21.31 locus; it has been hypothesized that its propensity for CNVs is responsible for the evolutionary toggling of this region that resulted in the distinct H1 and H2 haplotypes^34^. Due to the complex structural variation surrounding *LRRC37A* and the presence of its paralog *LRRC37A2*, it is challenging to genotype or sequence this region of the genome. As such, our analyses have not been able to separate the contribution of each gene and have considered them together. As a consequence of the CNVs in this region, genotyping data across *LRRC37A* and *LRRC37A2* is of low confidence and quality, and as such is excluded from GWAS analyses, as is visible in any association plot. Therefore, the association between *LRRC37A/2* variants and any disease or phenotype has never been tested, and it is likely that additional variation within *LRRC37A/2* itself is contributing to its altered expression and function that may also impact PD risk. Alternatively, differential epigenetic modifications or chromatin accessibility and looping may also be contributing to variable *LRRC37A/2* expression between sub-haplotypes. Unfortunately due to the relatively low frequency of different H1 sub-haplotypes, we do not yet have the brain tissues or iPSC lines available to test this possibility.

Little is known about the function of *LRRC37A/2*, although increased copy number has been associated with an increased antibody response to an Anthrax vaccine^35^, and overexpression in HeLa cells appeared to promote the formation of filopodia^33^, indicating that *LRRC37A/2* may be involved in the immune and inflammatory response, as well as with cellular migration and synapse formation. Furthermore, assessment of the GWAS atlas (https://atlas.ctglab.nl/) reveals that several immune phenotypes are significantly associated with H1/H2 haplotypes across the 17q21.31 locus, thus indicating this region is important for the regulation of immunological function. Despite conducting our RNA-seq analysis of *LRRC37A/2* overexpression in HEK293T cells, we observe enrichment of pathways consistent with these data; we found that increased *LRRC37A/2* expression upregulated cellular migration and chemotaxis pathways, which are both essential mechanisms involved in wound healing and inflammation. Within these pathways we observe increased expression of pro-inflammatory cytokines *IL17* and *IL32*, as well as the chemoattractant *IL16*, each of which are involved in the astrocytic inflammatory response^30, 31^.

Neuroinflammation of the substantia nigra is considered a characteristic feature of PD in addition to neuronal loss^36, 37^, and many genes associated with PD, such as *GBA*, *LRRK2* and *PINK1* are thought to have a role in the inflammatory response in astrocytes^24, 38^. Furthermore, the most significantly enriched pathway in our analysis, *Neuroactive-ligand receptor interaction*, is also involved in the inflammatory response and has previously been associated with PD; this pathway was significantly enriched in a functional assessment of PD GWAS signals^39^, and is targeted by microRNAs that are upregulated in a *Drosophila* model of PD^40^. We also observe upregulation of *TGFA*, the infusion of which into the forebrain of a rat model of PD increased the proliferation of neuronal and glial progenitors and the production of dopaminergic neurons to the substantia nigra^41^, indicating that this may be a protective growth factor against neuronal loss in PD. *LRRC37A/2* overexpression in HEK293T cells was therefore able to recapitulate pathways associated with PD, despite being a cell type with limited relevance to PD pathogenesis.

As these expression data were indicative of pathways relevant to astrocyte biology, and we found that *LRRC37A/2*-associated gene expression changes were apparent in iPSC-astrocytes but not iPSC-neurons, we hypothesized that this was likely the most relevant cell type for *LRRC37A/2* expression. Indeed, in human substantia nigra tissue we observe localization of LRRC37A/2 specifically in astrocytes. The contribution of astrocytic dysfunction to PD pathogenesis has gained attention in recent years, and is hypothesized to be a causal mechanism for the initiation and progression of PD^22–25^. Many genes associated with PD risk are expressed in astrocytes, the functions of which converge on the inflammatory response, lipid handling, mitochondrial health and lysosomal function^24^. Our finding that *LRRC37A/2* is expressed in astrocytes and plays a role in the inflammatory response is therefore consistent with known pathogenic mechanisms of PD.

The role of astroglial inflammation in PD is unclear; it has been reported as being both absent and severe in PD substantia nigra^22^. As a further complication, *in vitro* studies of human astrocyte cultures indicate that α-synuclein induces the release of pro-inflammatory α cytokines^22, 25^, but these cells also release protective molecules such as GDNF in response to dopaminergic neuronal damage, and such trophic support may benefit neuronal survival^22^. In addition, the substantia nigra is considered to be particularly susceptible in PD as dopaminergic neurons in this region are surrounded by the lowest proportion of astrocytes in the brain^42^. Whether an inflammatory response in this context would be protective or exacerbate neuronal death is therefore unknown. As our data suggest increased *LRRC37A/2* expression is protective and associated with increased expression of pro-inflammatory cytokines, astroglial inflammation in response to α-synuclein may therefore be protective. However, our use of HEK293T cells α limits our interpretation of these data, and further investigation of *LRRC37A/2* function in astrocytes is required.

Interestingly, we observe an interaction between LRRC37A/2 and soluble α-synuclein, as α well as co-localization of LRRC37A/2 with Lewy bodies in PD substantia nigra. The function and mechanism of this interaction is untested, although it is likely that a complex is formed in astrocytes and propagated to neurons. iPSC-astrocytes have been reported as expressing low levels of endogenous α-synuclein, which is increased in cells derived from PD patients^25^, and synuclein released from neuronal axon terminals is taken up by astrocytes^28^, which can be further transferred to neurons^23, 29^. This raises the possibility that LRRC37A/2 may influence release, aggregation and/or propagation.

In conclusion, we have identified novel sub-haplotype variants of the 17q21.31 H1 clade significantly associated with protection against PD. While the genetic association across this α-synuclein locus is typically ascribed to *MAPT* or *KANSL1*, we find evidence for the involvement of a novel gene, *LRRC37A/2,* in PD risk. We propose that in a similar mechanism to other PD-associated genes, *LRRC37A/2* is expressed in astrocytes and plays a role in the regulation of astroglial inflammation, specifically in the release of pro-inflammatory cytokines, chemotaxis and cellular migration. Importantly, we demonstrate that LRRC37A/2 interacts and co-localizes with - αsynuclein and Lewy bodies, thus indicating a potential modifying role in the formation of PD pathology. These findings link the genetic association at the 17q21.31 H1 locus with PD pathology, and support the hypothesis of astroglial dysfunction as a key contributing factor to PD disease pathogenesis.

## Methods

### Genotype data treatment

Case and control data from several cohorts from the International Parkinson’s Disease Genetics Consortium (IPDGC; NIA, GER, FIN, NL, SP, McGill)^6, 9, 10^ was kindly shared by Drs. Nalls, Singleton and Bandres-Ciga (NIH, Bethesda, MD; Tables S2, S4).

### Pre-imputation QC

Each dataset was obtained with different QC filters already applied, and so were all subsequently passed through the same, more stringent QC pipeline to ensure consistency. Plink v1.9^43^ was used to perform quality control for all datasets. First, SNPs were filtered by a 98% call rate, and remaining SNPs with a MAF < 1% were excluded. Individuals with < 98% genotyping rate were then removed. To determine and correct for population stratification, principal components analysis was carried out in combination with Hapmap YRI, CEU and CHB populations^44^ using EIGENSOFT^45^. Samples that did not cluster with the CEU European ancestry population were excluded. Identity by descent analysis was then conducted, and related individuals or potential sample duplicates (Z0 ≤ 0.8) were removed. We were unable to assess discordant sex information on data acquired from other sources, as the required information for this analysis was not provided to us. Variants that deviated from Hardy Weinberg equilibrium at a significance threshold < 1x10^-4^ were then removed. Chromosome 17 was then isolated and screened for strand mismatches.

### Imputation and post-imputation QC

Filtered chromosome 17 data from each cohort was submitted individually to the Michigan imputation server^46^ (https://imputationserver.sph.umich.edu) and imputed against the HRC r1.1 2016 panel using Eagle v2.3 phasing. Following imputation, SNPs with an r2 < 0.3 were removed, and remaining SNPs were filtered for a 99% call rate. Genotyping call rates for individuals were again filtered at 99%, and SNPs that deviated from Hardy-Weinberg equilibrium at a significance threshold < 1x10^-6^ were excluded. Individual cohorts were then merged, and finally filtered once more with a SNP call rate of 99%. Prior to analysis, variants were filtered to exclude SNPs with a MAF < 0.01.

### Single SNP association analyses

Logistic regression association analyses using an additive model were carried out in SNP and Variation Suite v8.8.1 (SVS8) software (Golden Helix, Inc., Bozeman, MT, www.goldenhelix.com). As all potential covariate information was not available, the model was corrected using the first 10 principal components as calculated by SVS8. Associations were initially carried out on the entire cohort in order to confirm the 17q21.31 H1/H2 haplotype association. The data was then filtered for H1 homozygotes only, using tag SNP rs8070723 and the association analysis was repeated with the same parameters.

Meta-analysis of SNP effects across multiple datasets was carried out using the R package rmeta^47^ using both Random Effects (DerSimonian-Laird) and Fixed Effects (Mantel-Haenszel) approaches. Calculation and visualization of linkage disequilibrium (LD) over large genomic ranges was carried out in SVS8 using both r2 and D’. Inspection of LD between individual SNPs of interest was carried out using Haploview^48^.

### Haplotype block construction and association

Haplotype blocks were constructed in SVS8 using the D’ measure of LD. Blocks were defined using guidelines as described by Gabriel et al (2002)^49^. Each block contained a maximum of 15 markers within 160kb of each other, with a D’ upper confidence bound ≥ 0.98 and a lower confidence bound ≥ 0.7. Haplotypes were estimated using an expectation-maximization (EM) algorithm with 50 iterations, and a convergence tolerance of 0.0001. Sub-haplotypes with a frequency < 0.01 were excluded from further analysis. Case-control association analyses were carried out per block using a logistic regression model. Odds ratios and associated Fisher’s exact p-values were calculated for each sub-haplotype within each block using the R package epitools^50^.

### Human brain expression analysis

Publicly available RNA-seq expression data from human postmortem dorsolateral prefrontal (PFC) and temporal (TCX) cortices (Table S3) and associated genotype data were obtained from Synapse (synapse.org; The Religious Orders Study and Memory and Aging Project (ROSMAP) syn3219045; MayoRNAseq syn5550404; CommonMind Consortium syn2759792). Genotype data for chromosome 17 underwent the same QC and imputation pipeline as described above. Data were stratified by 17q21.31 H1/H2 haplotype using the H2 tag SNP rs8070723. For sub-haplotype analysis, blocks previously defined in the PD analysis were applied to the genotype data and haplotypes were estimated in the same manner. Statistical analysis was carried out in R version 3.4.0. For analysis of *MAPT* splicing, percent spliced in (PSI) values were generated for exons 2, 3 and 10 using Mixture of Isoforms (MISO)^51^. Gene expression and PSI residuals were generated by linear regression using sex, age of death, post-mortem interval and RNA integrity score as covariates. The resulting residuals were then transformed into z-scores and combined across datasets. Statistical differences in gene expression between genotypes and sub-haplotypes were determined by linear regression applied to the adjusted and combined z-scores.

### dPCR

Human genomic DNA and accompanying genotype data was kindly provided by Drs. Raj, Crary and Charney (Mount Sinai School of Medicine, NY) and by the Alzheimer’s Disease Research Center (ADRC; Table S3). Sub-haplotypes were called from these genotype data in the same manner as described above. To examine copy number variation in the 17q21.31 locus, digital PCR was carried out using the ThermoFisher QuantStudio 3D digital PCR chip system. Taqman dPCR probes for loci within the alpha, beta and gamma CNV regions^20^, as well as within *LRRC37A* and *MAPT* were selected for analysis (Table S5).

### Cell lines

Human induced pluripotent stem cells (iPSCs) were obtained from the Knight Alzheimer’s Disease Research Center at Washington University^52^, the NIH Childhood-onset Schizophrenia study^53^, and the UCI ADRC (Table S6). The Icahn School of Medicine at Mount Sinai IRB reviewed the relevant operating protocols as well as this specific study and determined it was exempt from approval.

### Cell culture

Unless otherwise specified, all cell culture materials were obtained from ThermoFisher Scientific. Human embryonic kidney cells (HEK293T) were cultured in Dulbecco’s Modified Eagle Medium/F-12 with HEPES, supplemented with 10% fetal bovine serum (FBS) and 1% Penicillin-Streptomycin. Cells were passaged every 3-4 days using Trypsin-EDTA (0.25%). For *LRRC37A*/2 overexpression experiments, HEK293T cells were seeded at a density of 1.4x10^5^ cells per well in 6-well plates and transfected with 0.5-2.5ug of *LRRC37A* plasmid (Origene) or empty vector control (Origene) using Lipofectamine 3000. Cells were harvested 48 hours after transfection.

For qRTPCR and protein biochemistry experiments, iPSC lines (Table S6) were maintained in complete StemFlex media supplemented with 1% penicillin/streptomycin on Matrigel (BD biosciences), and were differentiated to neural progenitor cells (NPCs) as previously described^54^. Forebrain neuron-enriched cultures and astrocyte cultures were differentiated from NPCs as previously described^54, 55^. Neuronal and astrocytic identity was confirmed by immunofluorescence for common neuronal and astrocytic markers (MAP2 (Abcam), Tuj1 (Cell Signaling Technologies), S100β (Sigma Aldrich) and EAAT1 (Abcam).

Genomic DNA was extracted using the DNeasy Blood and Tissue kit (Qiagen) and underwent genotyping with Taqman assays for H2 tag SNPs rs8070723 and rs1052553 in order to confirm 17q21.31 haplotype.

### qRT-PCR

RNA was extracted from HEK293T cells, iPSC-derived neurons, astrocytes and human brain tissue using the RNeasy Mini kit (Qiagen) and reverse transcribed using the High-Capacity RNA-to-cDNA kit (ThermoFisher Scientific). Gene expression was measured by commercially available Taqman qRTPCR assays.

### RNA-seq

RNA was prepared as described above. Library preparation with poly-A selection and sequencing with 150 base pair paired-end reads was carried out at Genewiz. Sequenced reads were trimmed for Illumina TruSeq adapters, and quantified for gene expression values in TPM (Transcripts Per Kilobase Million) using Salmon^56^ guided by the GENCODE human transcriptome model (GRCh38 version 28, Ensembl 92). TPM data was imported into the R (version 3.5.1) programming environment for visualization and analysis, and differential expression of *LRRC37A*/2 overexpression compared to the control was analyzed using the moderated t-test implemented in limma^57^. Differentially expressed genes (DEGs) were defined < 0.05. Gene set enrichment analysis was performed with the Broad Institute’s MSigDB annotations^58^. Analysis of GO enrichment terms was carried out using g:Profiler (https://biit.cs.ut.ee/gprofiler/gost)^59,60^ and visualized using Cytoscape v3.7.1^61^ with the EnrichmentMap^62^ plugin. Additional pathway analyses were carried out using Ingenuity Pathway Analysis (QIAGEN Inc., https://www.qiagenbioinformatics.com/products/ingenuitypathway-analysis) using genes with a fold change ± ≥ 1.

### Protein biochemistry

Membrane and cytosolic proteins were isolated from HEK293T cells, iPSC-derived neurons and iPSC-derived astrocytes using the MEM-PER Plus Membrane Protein Extraction Kit (ThermoFisher Scientific), and protein concentrations were determined by bicinchoninic acid (BCA) assay (ThermoFisher Scientific). For western blotting, protein fractions were subject to SDS-PAGE electrophoresis through BOLT Bis-Tris gels (ThermoFisher Scientific) and were blotted onto nitrocellulose membranes. Membrane fractions were confirmed by labelling with an anti-pan-Cadherin antibody (Cell Signaling Technology), and cytosolic fractions were confirmed by labelling with anti-HSP90 (Cell Signaling Technology). Membranes were stripped using Restore plus western blot stripping buffer and re-probed with an anti-LRRC37A antibody (ThermoFisher Scientific).

### OPAL multiplex labelling

Formalin fixed paraffin embedded substantia nigra sections from human controls (N=4), PSP (N=5) and PD (N=5) cases were acquired from the Mount Sinai Neuropathology Brain Bank, with neuropathological diagnosis being determined by Dr. John Crary. All post-mortem tissues were collected in accordance with the relevant guidelines and regulations at the Icahn School of Medicine at Mount Sinai. Multiplexed immunofluorescent staining was carried out on 4-6µm sections using the Opal Polaris 7 color IHC detection kit (Akoya biosciences) according to manufacturer’s instructions. Briefly, slides were baked for 1 hour at 65°C, then deparaffinized with xylene and rehydrated with a graded series of ethanol concentrations. For epitope retrieval, slides were microwaved in AR buffer for 45s at 100% power, followed by an additional 15 minutes at 20% power. After cooling, slides were blocked for 10 minutes in blocking buffer then incubated with the first primary antibody at room temperature for 30 minutes. Slides were rinsed three times in TBS-T, then incubated with the secondary polymer HRP for 1 hour at room temperature. After additional washes, the first Opal fluorophore was incubated with the slides for 10 minutes at room temperature, followed by further washes in TBS-T. This process was repeated from the microwave treatment step for each additional primary antibody, followed by one final repetition of the microwave treatment to strip the primary-secondary antibody complex from the tissue. Once all primary antibodies had been introduced, slides were counterstained with DAPI for 5 minutes at room temperature, washed with TBS-T and coverslips were mounted using ProLong Diamond Antifade mounting reagent (ThermoFisher Scientific). Multispectral imaging was carried out using the Vectra Quantitative Pathology Imaging system, applying quantitative unmixing of fluorophores and removal of tissue autofluorescence. Images were visualized using the HALO image analysis platform (Indica Labs).

### Co-Immunoprecipitation

Frozen substantia nigra tissue was selected from the same Control (N=3), PSP (N=3) and PD (N=3) cases used for OPAL multiplex immunofluorescence. Protein lysates were generated using cell lysis buffer (NEB) and brief sonication on ice, followed by centrifugation to pellet insoluble material. Co-immunoprecipitation was carried out using the Dynabeads Protein G immunoprecipitation kit (ThermoFisher Scientific), with an anti-α-synuclein antibody (Abcam) α as bait. Proteins bound to beads were eluted and assayed by western blot (as described above) and probed with an anti-LRRC37A antibody (ThermoFisher Scientific). Whole protein lysate and IgG only controls were run on the same blots.

## DATA SHARING

All aligned read counts and FASTQ files for *LRRC37A*-overexpressing HEK293T cells will be deposited to the Gene Expression Omnibus once the manuscript is accepted for publication.

## ACKNOWLEDGEMENTS

This work was supported by funding from the BrightFocus Foundation (KRB), Association for Frontotemporal Degeneration (KRB) and CurePSP (KRB). The recruitment and clinical characterization of research participants at Washington University were supported by NIH P50 AG05681, P01 AG03991, and P01 AG026276. NIH AG062683 (J.TCW), Rainwater Charitable Organization (CMK), NIH AG046374 (CMK). The McGill cohort was supported by grants from the Michael J. Fox Foundation, the Canadian Consortium on Neurodegeneration in Aging (CCNA), the Canada First Research Excellence Fund (CFREF), awarded to McGill University for the Healthy Brains for Healthy Lives (HBHL) program, and Parkinson Canada. ZGO is supported by the Fonds de recherche du Québec - Santé (FRQS) Chercheurs-boursiers award given with Parkinson Quebec, and is a Parkinson Canada New Investigator awardee. The access to the McGill participants for this research has been made possible in part thanks to the Quebec Parkinson’s Network (http://rpq-qpn.ca/en/). NIH [R01AG054008, R01NS095252, R01AG060961, R01 AG060961] to JFC, the Rainwater Charitable Foundation/Tau Consortium (JFC), Genentech/Roche, and an Alexander Saint-Amand Scholarship (JFC). This work was supported in part by the Intramural Research Programs of the National Institute of Neurological Disorders and Stroke (NINDS), the National Institute on Aging (NIA), and the National Institute of Environmental Health Sciences both part of the National Institutes of Health, Department of Health and Human Services: project numbers 1ZIA-NS003154, Z01-AG000949-02 and Z01- ES101986.

## AUTHOR CONTRIBUTIONS

*Conceptualization:* KRB, JFC, AMG. *Methodology*: KRB, JDC. *Validation:* KRB, DAP. *Formal Analysis:* KRB, DAP, YL, SB-C, ZG-O, PH, AS, SB. *Investigation*: KRB, DAP, YL. *Resources:* AC, CMK, SJF, BHK, IP, YJP, AC, TR, JFC, AMG. *Data Curation*: KRB, YL, SB-C, ZG-O, PH, AS, SB, IPDGC, *Writing – Original Draft:* KRB. *Writing – Review and Editing*: KRB, YL, AER, SB-C, ZG-O, CMK, IP, TR, JFC, AMG. *Visualization:* KRB. *Supervision*: AMG. *Funding Acquisition:* KRB, JFC, AMG

## SUPPLEMENTARY FIGURES LEGENDS

**Figure S1.**
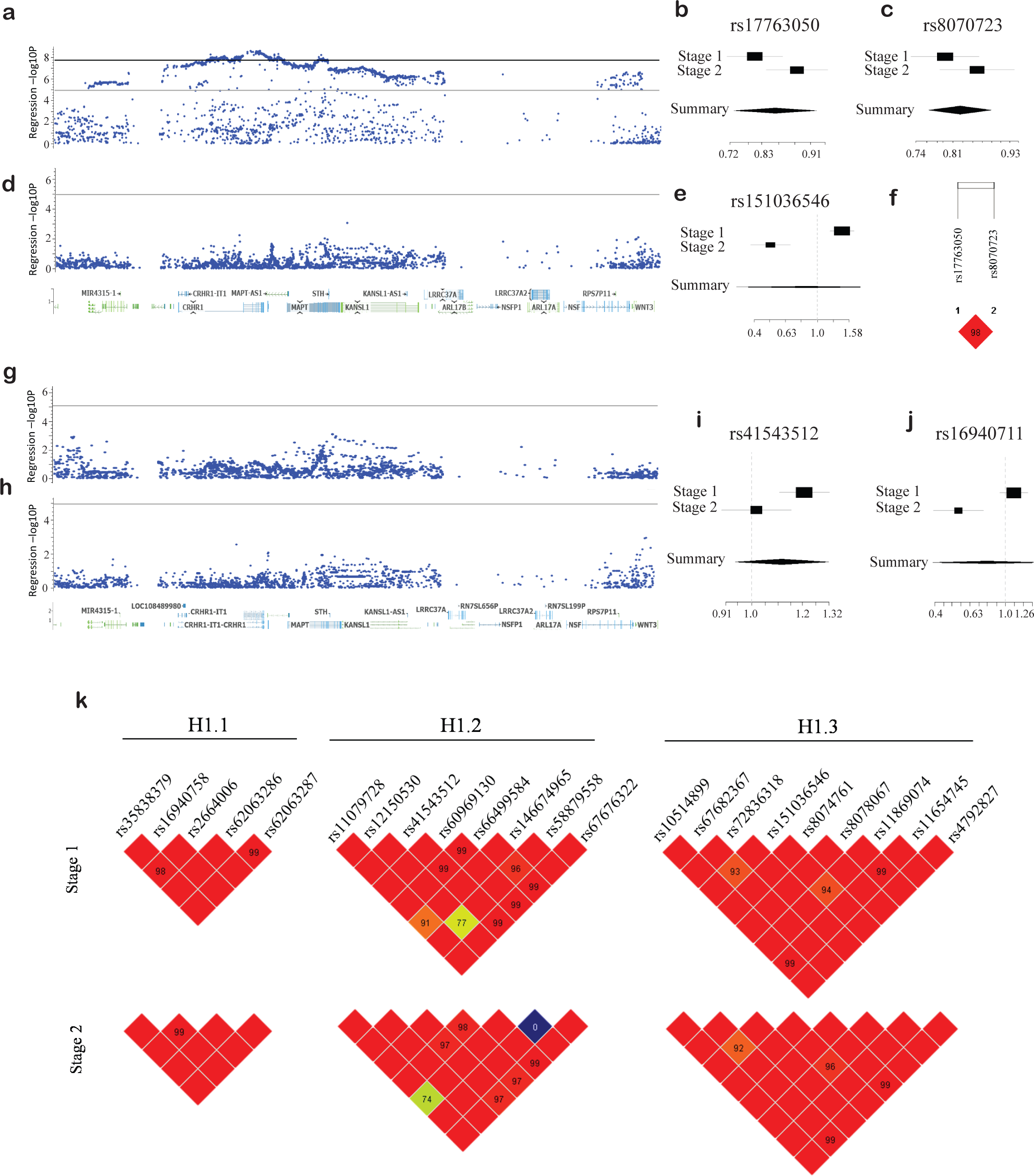
The 17q21.31 locus is genetically associated with PD. Related to Figure 1. **A.** –log_10_ regression *p*-values of SNP associations with PD in Stage 1 data spanning the 17q21.31 locus. Grey line indicates genome-wide suggestive *p*-value of 1x10^-5^, and black line indicates genome-wide significant *p*-value of 5x10^-8^ **B-C.** Odds ratios (ORs) and random effects meta-analysis (Summary) with 95% confidence intervals for ***B.*** the top association SNP from Stage 1 data and **C.** H2 tag SNP rs8070723 in Stage 1 and Stage 2. **D.** –log_10_ regression *p*-values of SNP associations with PD in Stage 2 data spanning the 17q21.31 locus. Grey line indicates genome-wide suggestive *p*-value of 1x10-5 **E.** ORs and random effects meta-analysis (Summary) with 95% confidence intervals for the top association SNP from Stage 2 data **F.** D’ LD between the Stage 1 top association SNP and H1/H2 tag SNP rs8070723 **G-H.** –log_10_ regression *p-*values of SNP associations with PD in H1 homozygotes in ***G.*** Stage 1 and ***H.*** Stage 2 data spanning the 17q21.31 locus. Grey line indicates genome-wide suggestive *p*- value of 1x10^-5^. **I-J.** ORs and random effects meta-analysis (Summary) with 95% confidence intervals for top SNP associations in ***I.*** Stage 1 and ***J.*** Stage 2 PD data. **K.** LD (D’) between SNPs within each significant PD-associated sub-haplotype block; H1.1 (chr17:44040184-44041992), H1.2 (chr17:44090196-44097249) and H1.3 (chr17:44119987- 44131305) in Stage 1 (top) and Stage 2 (bottom) data. Red indicates high LD; diamonds with no value denote D’ = 1. D’ values <1 are indicated for each relevant SNP pair.

**Figure S2.**
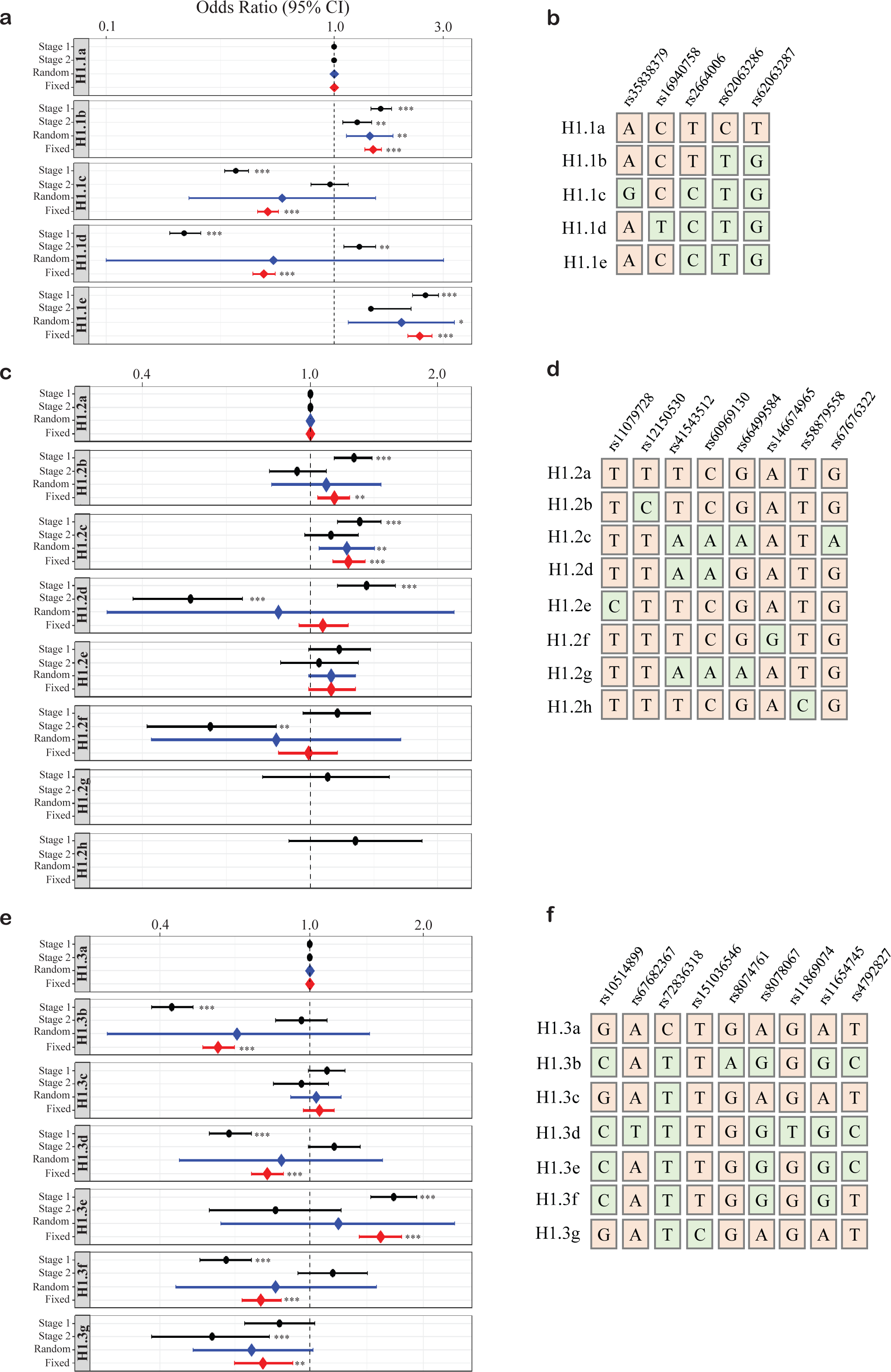
Sub-haplotypes within H1 sub-haplotype blocks are associated with variable risk for PD. Related to Figures 1 and 2. **A-B.** Block H1.1 ***A***. Sub-haplotype SNP genotypes (green squares denote minor allele variation relative to the most common haplotype) and ***B.*** related ORs and 95% confidence intervals in Stage 1 and Stage 2 data (black), random effects meta-analysis (blue) and fixed effects meta-analysis (red). **C-D.** Block H1.2 ***C.*** Sub-haplotype SNP genotypes (green squares denote minor allele variation relative to the most common haplotype) and ***D.*** related ORs and 95% confidence intervals in Stage 1 and Stage 2 data (black), random effects meta-analysis (blue) and fixed effects meta-analysis (red). **E-F**. Block H1.3 ***E.*** Sub-haplotype SNP genotypes (green squares denote minor allele variation relative to the most common haplotype) and ***F.*** related ORs and 95% confidence intervals in Stage 1 and Stage 2 data (black), random effects meta-analysis (blue) and fixed effects meta-analysis (red). \**p* < 0.05, \*\**p* < 0.01, \*\*\**p* < 0.001

**Figure S3.**
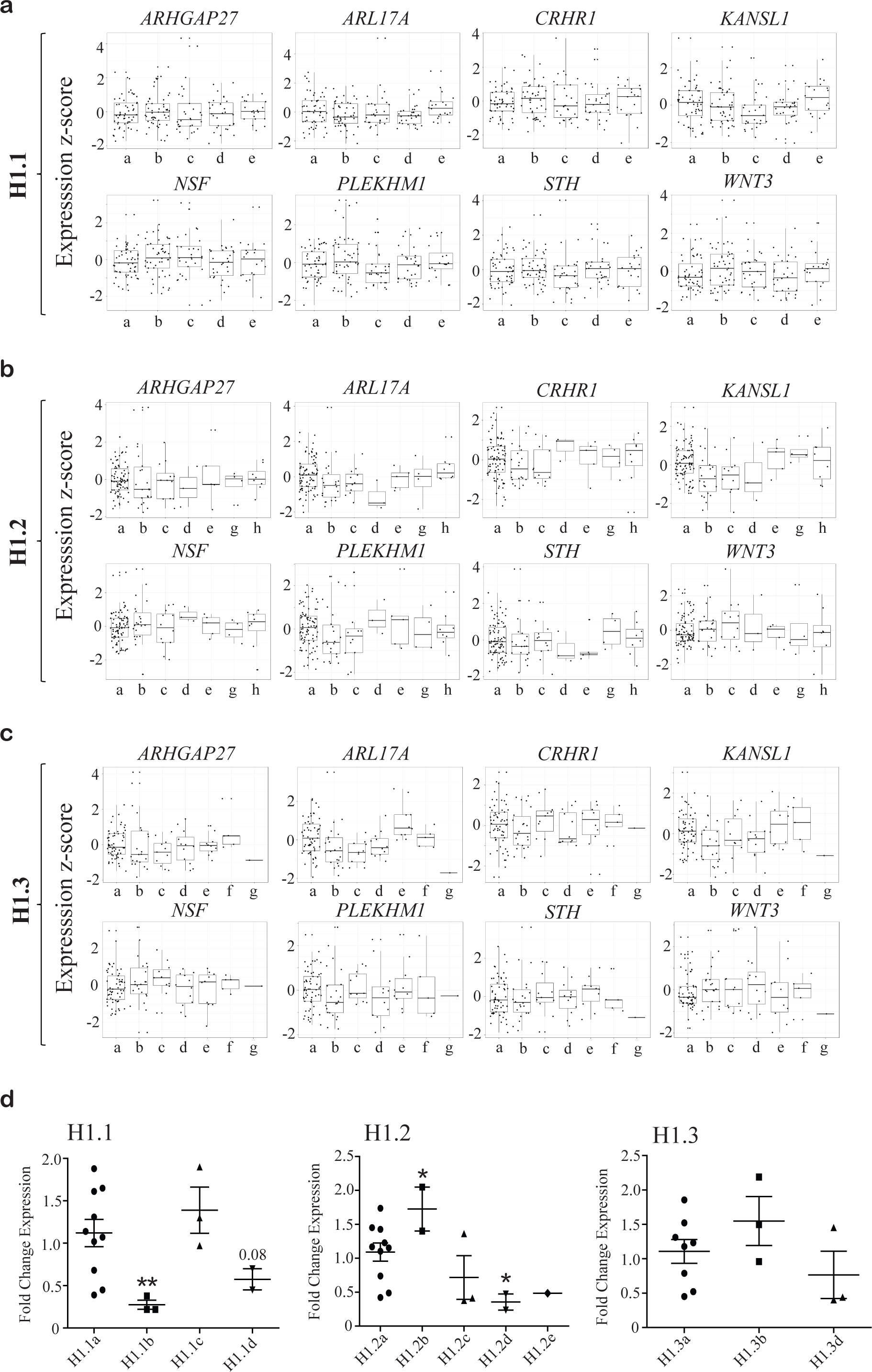
PD-associated sub-haplotype blocks are not associated with the expression of most genes within the 17q21.31 locus. Related to Figure 2. **A-C.** Combined residual Z-scores for expression of genes across the 17q21.31 locus for individuals homozygous for each sub-haplotype in blocks ***A.*** H1.1, ***B.*** H1.2 and ***C.*** H1.3 human brain RNA-seq data. **D.** qRTPCR for *LRRC37A* in human PFC brain tissue for sub-haplotypes across blocks H1.1, H1.2 and H1.3. N = 1-10, * *p*<0.05.

**Figure S4.**
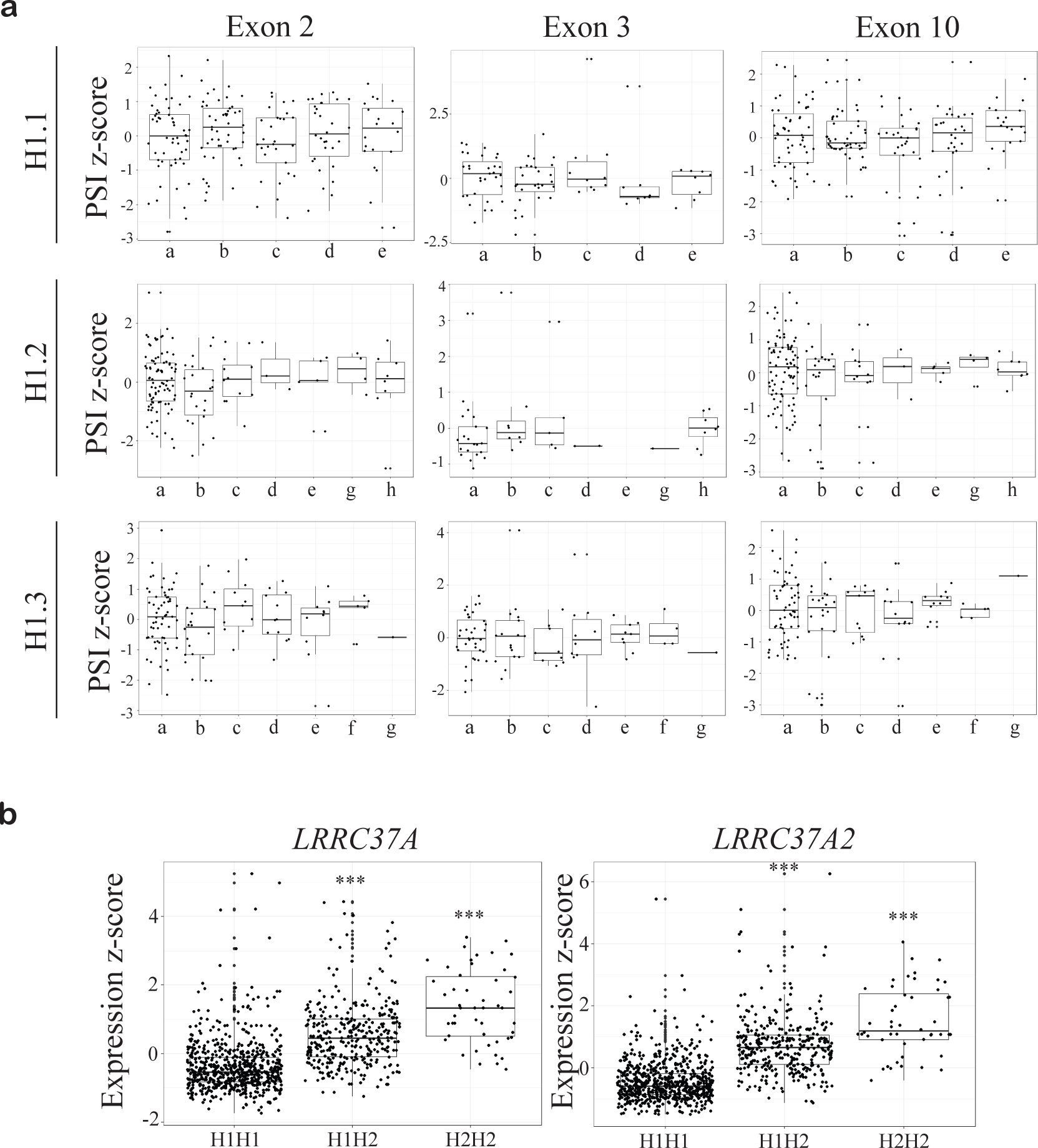
PD-associated sub-haplotype blocks are not associated with *MAPT* splicing. Related to Figure 2. **A.** Combined residual Z-scores for *MAPT* exons 2, 3 and 10 percent spliced in (PSI) for individuals homozygous for each sub-haplotype in blocks H1.1, H1.2 and H1.3 in human brain RNA-seq data. **B.** Combined residual Z-scores for *LRRC37A* and *LRRC37A2* expression in 17q21.31 H1 and H2 haplotype clades in human brain RNA-seq data. All statistical comparisons are against the most common allele. \**p* < 0.05, \*\**p* < 0.01, \*\*\**p* < 0.001.

**Figure S5.**
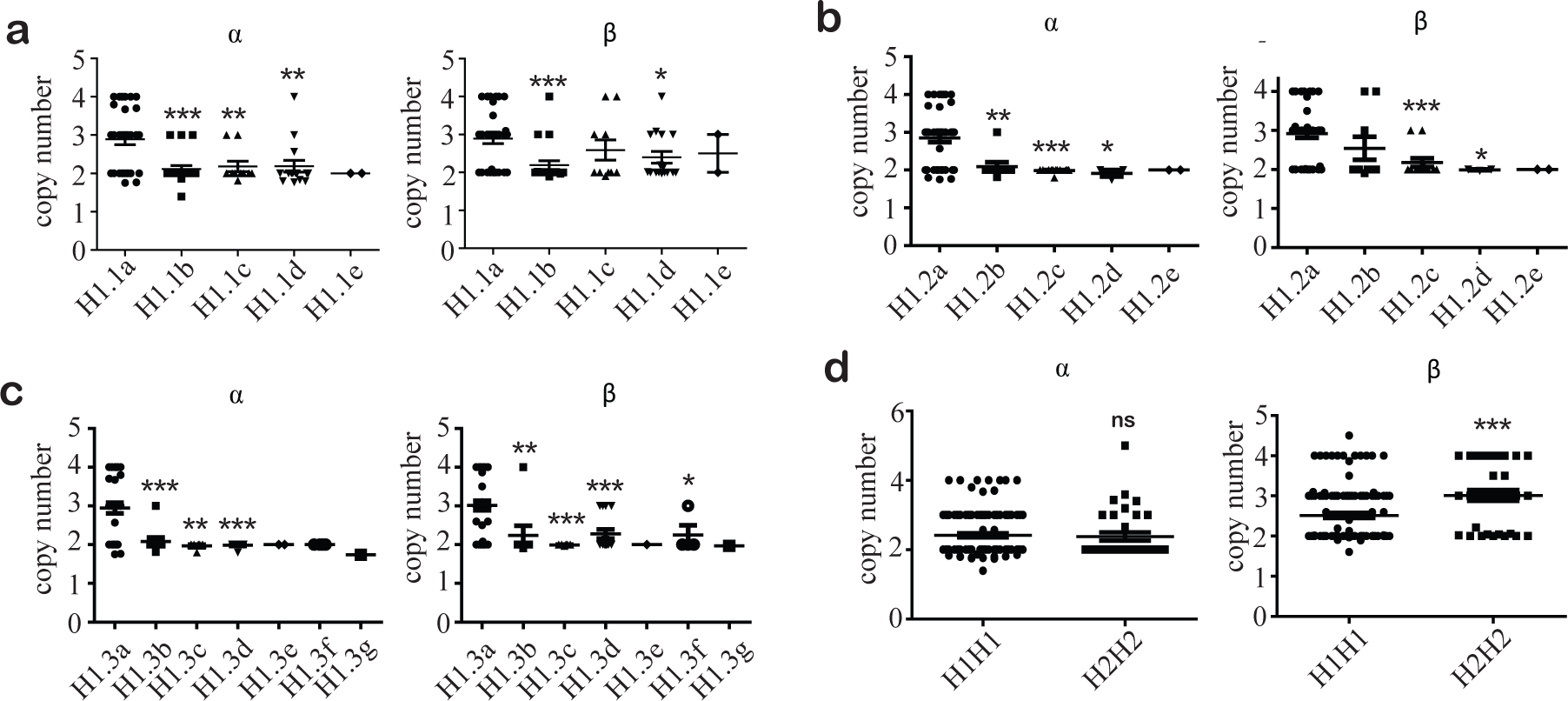
Alpha and beta regions have more variable copy numbers in the most common block sub-haplotypes, but do not vary with PD risk. Related to Figure 2. **A-C.** Copy number of alpha and beta regions across sub-haplotypes in blocks ***A***. H1.1, ***B.*** H1.2 and ***C.*** H1.3. **D.** Copy number of alpha and beta regions between H1 and H2 homozygotes. All statistical comparisons are against the most common sub-haplotype. ns = not significant, \**p* < 0.05, \*\**p* < 0.01, \*\*\**p* < 0.001.

**Figure S6.**
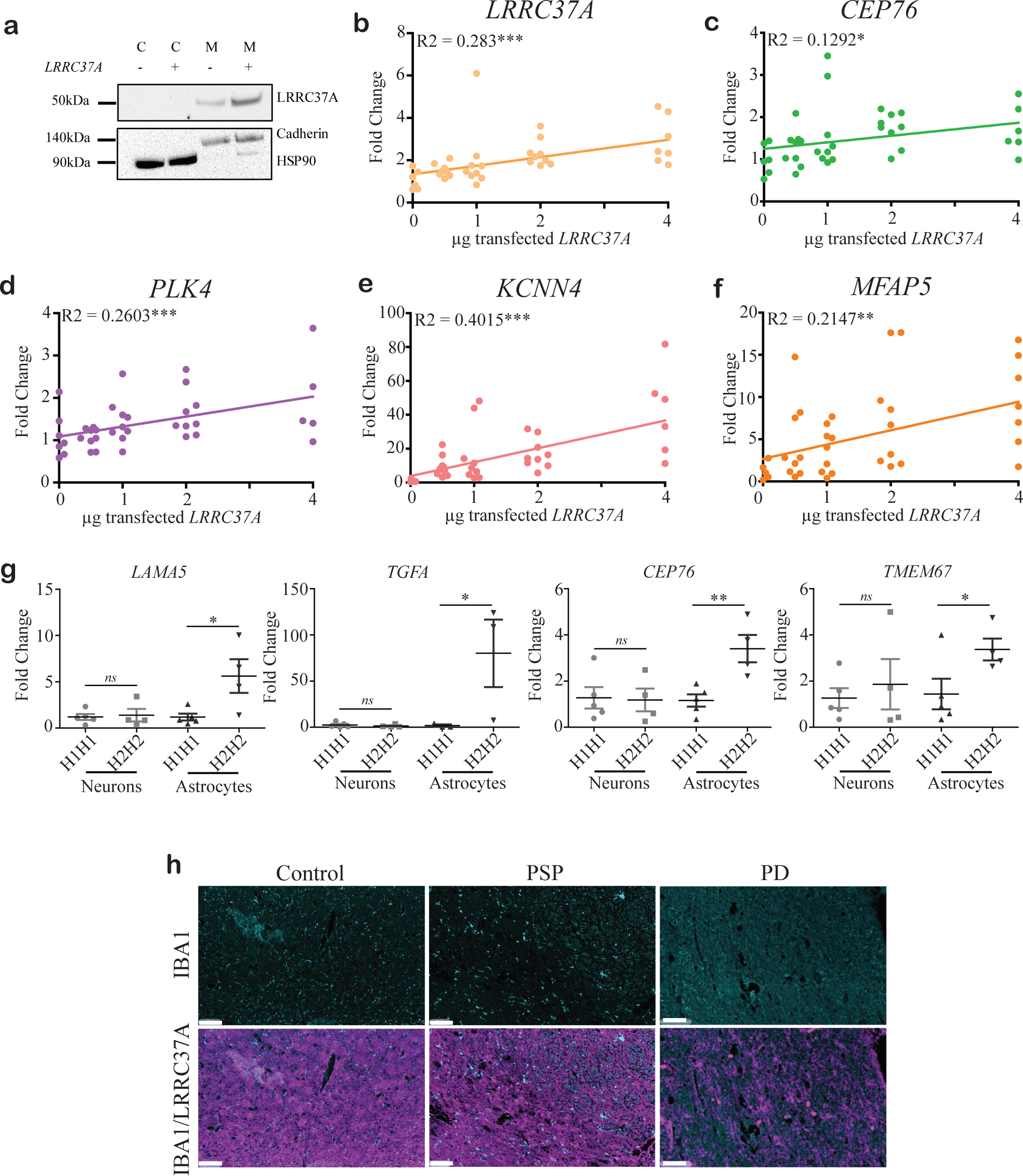
LRRC37A/2 is a membrane-associated protein with the greatest effect in astrocytes, but is not expressed in microglia. Related to Figures 3 and 4. **A.** Western blots for LRRC37A in cytosolic (C) and membrane (M) fractions from HEK293T cells overexpressing *LRRC37A/2* or an empty vector control. Cytosolic fractions were confirmed by labelling with an anti-HSP90 antibody, and membrane fractions were confirmed by labelling with an anti-Pan-Cadherin antibody, N=3. **B-F.** qRT-PCR fold change expression of differentially expressed genes ***B.*** *LRRC37A*, ***C.****CEP76*, ***D.*** *PLK4*, ***E.*** *KCNN4* and ***F***. *MFAP5* in *LRRC37A/2*-overexpressing cells from RNA-seq data, following titration of *LRRC37A/2* overexpression in HEK293T cells. N=9, \**p*<0.05, \*\**p*<0.01, \*\*\**p*<0.001. **G.** qRT-PCR fold change expression of differentially expressed genes following *LRRC37A/2* overexpression in H1 and H2 homozygote iPSC-derived neurons and astrocytes, N=3 in duplicate, \**p*<0.05, \*\**p*<0.01, \*\*\**p*<0.001. **H.** Representative images demonstrating no co-localization of microglia (IBA1) and LRRC37A/2 in control, PSP and PD substantia nigra. N=4-5. Scale bar = 100µm.

## SUPPLEMENTARY TABLES

**Table S1.**
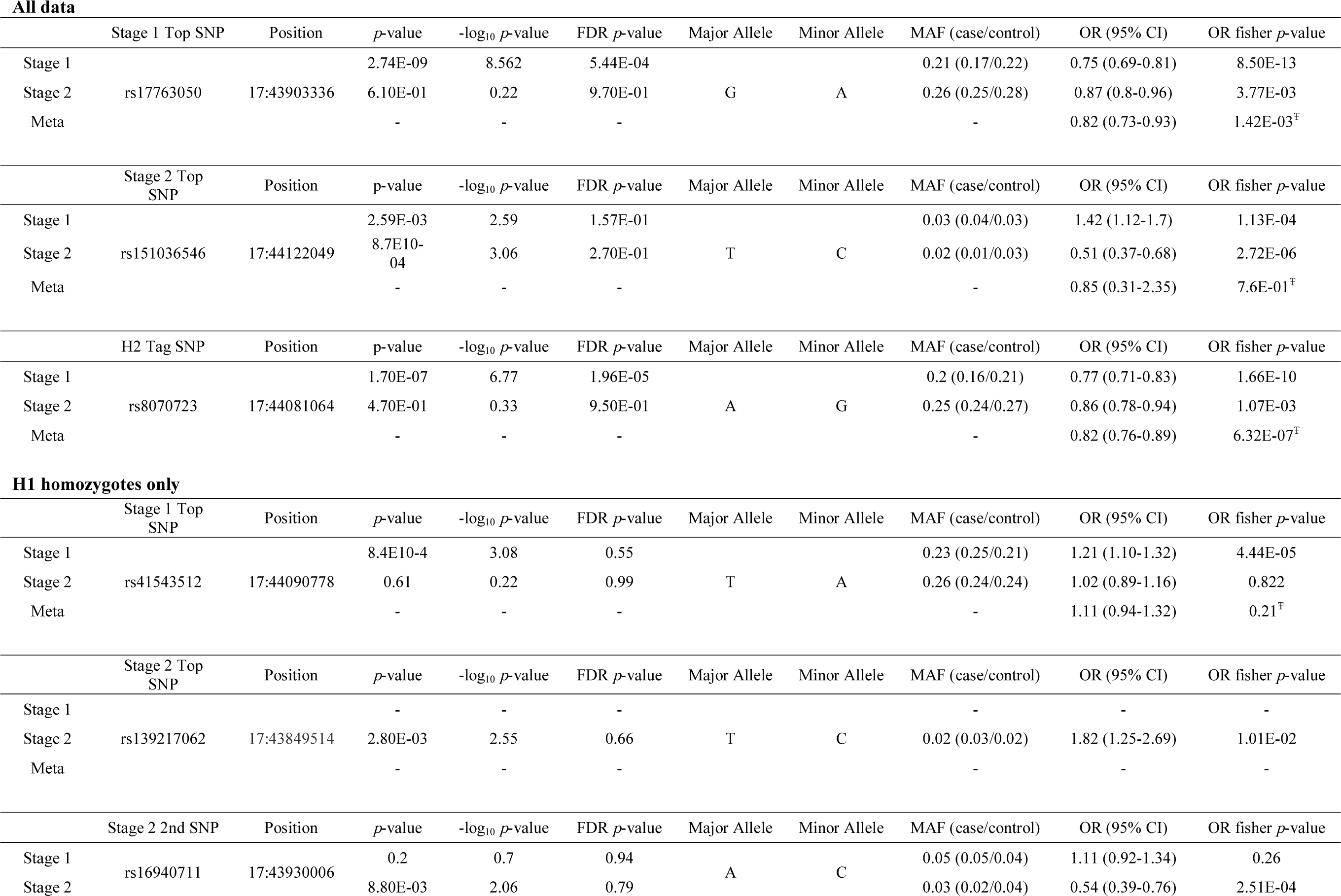
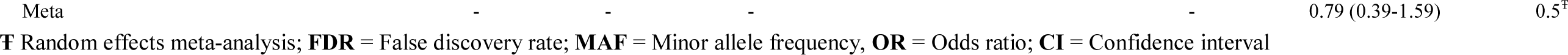
Top SNPs in the 17q21.31 locus associated with PD in complete data and H1 homozygote analyses

**Table S2.** Data summary for Stage 1 and Stage 2 PD analyses

|  | **Cohort** | **Array** | **Total N**<br>**(Case/Control)** | **H1H1**<br>**(Case/Control)** | **H2H2**<br>**(Case/Control)** | **#SNPs used for**<br>**imputation**<br>**(Chr17)** | **#Imputed SNPs**<br>**(Chr17) R2 >= 0.3** |
| --- | --- | --- | --- | --- | --- | --- | --- |
| Stage 1 | NIA (IPDGC) | Illumina 550K v1 & v3 | 3906 (905/3001) | 2447 (606/1841) | 161 (31/130) | 11941 | 634666 |
|  | NL (IPDGC) | Illumina 550K & 610K | 2725 (767/1958) | 1654 (500/1154) | 144 (39/105) | 13457 | 530635 |
|  | FIN (IPDGC) | Illumina 370K | 860 (368/492) | 732 (316/416) | 7 (4/3) | 7970 | 357255 |
|  | GER (IPDGC) | Illumina 550K v1 | 1673 (740/933) | 1121 (527/594) | 51 (19/32) | 11803 | 481302 |
|  | Merged | - | 9164 (2780/6384) | 5932 (1937/3995) | 378 (94/284) | - | 295782 |
| Stage 2 | McGill (IPDGC) | Illumina Omini Express<br>& NeuroChip v.1.0 | 1485 (582/903) | 859 (354/505) | 70 (19/51) | 17467 | 515155 |
|  | SP (IPDGC) | NeuroChip v.1.0 & v.1.1 | 3444 (2117/1327) | 1861 (1184/677) | 243 (130/113) | 16590 | 493744 |
|  | Merged | - | 4929 (2699/2230) | 2925 (1538/1182) | 313 (149/164) | - | 368729 |

**Table S3.** Sample summary for RNA-seq expression analyses, dPCR copy number variation analyses and qRTPCR.

|  |  | Brain<br>Region/Tissue | N | #H1H1 | #Homozygote<br>block1 H1 sub-haplotypes | #Homozygote<br>block2 H1 sub-haplotypes | #Homozygote<br>block3 H1 sub-haplotypes | #H2H2 |
| --- | --- | --- | --- | --- | --- | --- | --- | --- |
| RNA-seq<br>Analysis | AMP-AD<br>ROSMAP | PFC | 450 | 289 | 64 | 61 | 56 | 19 |
|  | AMP-AD MAYO | TCX | 276 | 185 | 49 | 45 | 38 | 7 |
|  | CommonMind | PFC | 624 | 347 | 70 | 34 | 28 | 21 |
|  | Total | - | 1350 | 821 | 183 | 140 | 122 | 47 |
| dPCR | MSMD | Blood | 66 | 49 | 37 | 30 | 32 | 17 |
|  | ADRC | Blood | 33 | 21 | 16 | 9 | 13 | 12 |
|  | Charney LB | Blood | 20 | 17 | 13 | 12 | 9 | 3 |
|  | Charney PD | PFC | 41 | 34 | 18 | 18 | 15 | 4 |
|  | Total | - | 160 | 121 | 84 | 69 | 69 | 36 |
| qRT-PCR | Charney PD | PFC | 41 | 34 | 18 | 18 | 15 | 4 |

**PFC** = prefrontal cortex; **TCX** = temporal cortex; **AMP-AD** = Accelerating Medicines Partnership Alzheimer's disease; **ROSMAP** = Religious Orders Study and Memory and Aging Project; **MSMD** = Mount Sinai Movement Disorders; **ADRC** = Alzheimer's Disease Research Center; **LB** = Living brain; **PD** = Parkinson's disease

**Table S4.** Quality control summary for Stage 1 and Stage 2 PD data analysis

|  | Stage 1 |  |  |  |  | Stage 2 |  |  |
| --- | --- | --- | --- | --- | --- | --- | --- | --- |
|  | NIA | NL | GER | FIN | Total | McGILL | SPAIN | Total |
| Initial sample size | 4005 | 2796 | 1686 | 883 | 9370 | 1847 | 3444 | 5291 |
| Missingness | 37 | 3 | 10 | 21 | 71 | 5 | 0 | 5 |
| Relatedness | 6 | 68 | 3 | 2 | 79 | 0 | 0 | 0 |
| Ancestry | 56 | 0 | 0 | 0 | 56 | 0 | 0 | 0 |
| Missing phenotype | 0 | 0 | 0 | 0 | 0 | 357 | 0 | 357 |
| Excluded (n) | 99 | 71 | 13 | 23 | 206 | 362 | 0 | 362 |
| Excluded (%) | 2.47 | 2.54 | 0.77 | 2.60 | 2.20 | 19.60 | 0.00 | 19.60 |
| Final sample size | **3906** | **2725** | **1673** | **860** | **9164** | **1485** | **3444** | **4929** |
| Cases | 905 | 767 | 740 | 368 | 2780 | 582 | 2117 | 2699 |
| Controls | 3001 | 1958 | 933 | 492 | 6384 | 903 | 1327 | 2230 |

**Table S5.**
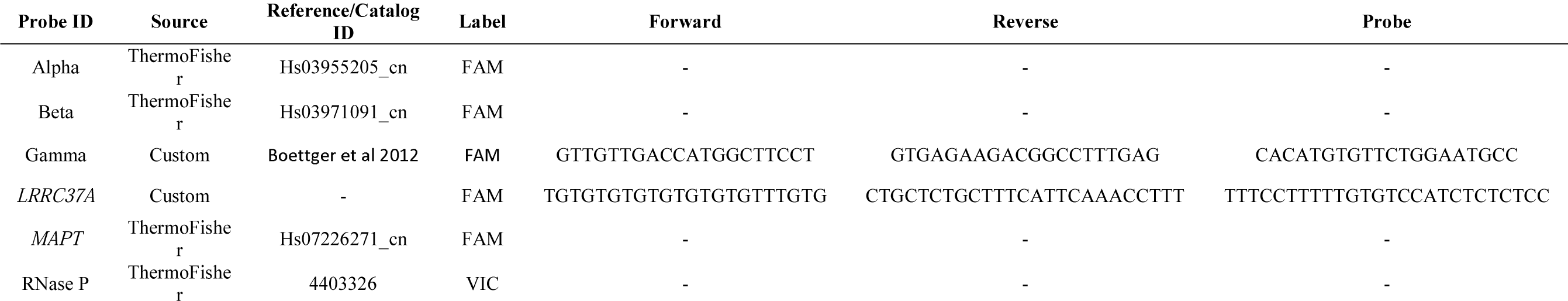
dPCR probe design

**Table S6.** Summary of iPSC line sources and 17q21.31 haplotypes.

| **Line ID** | **Source** | **Sex** | **Haplotype** |
| --- | --- | --- | --- |
| F11349 | ADRC | Male | H2H2 |
| F0510.2Δ2H1 | ADRC | Male | H1H1 |
| FA12455 | ADRC | Female | H2H2 |
| F13505 | ADRC | Female | H1H1 |
| 3182-3 | NIHC<br>Z | Female | H2H2 |
| F11421Δ2A07 | ADRC | Female | H1H1 |

**ADRC** = Knight Alzheimer's Disease Research Center at Washington University; **NIHCZ** = NIH Childhood-onset Schizophrenia study

## REFERENCES

1. Jun G, Ibrahim-Verbaas CA, Vronskaya M, Lambert J-C, Chung J, Naj AC. a Novel Alzheimer Disease Locus Located Near the. 2016; 21: 108–17.

2. Kouri N, Ross OA, Dombroski B, et al. Genome-wide association study of corticobasal degeneration identifies risk variants shared with progressive supranuclear palsy. Nat Commun 2015; 6: 1–7.

3. Höglinger GU, Melhem NM, Dickson DW, et al. Identification of common variants influencing risk of the tauopathy progressive supranuclear palsy. Nat Genet 2011; 43: 699–705.

4. Chen JA, Chen Z, Won H, et al. Joint genome-wide association study of progressive supranuclear palsy identifies novel susceptibility loci and genetic correlation to neurodegenerative diseases. Mol Neurodegener 2018; 13: 1–11.

5. Pastor P, Ezquerra M, Perez JC, et al. Novel Haplotypes in 17q21 Are Associated with Progressive Supranuclear Palsy. Ann Neurol 2004; 56: 249–58.

6. Bandrés-ciga S, Ryan T, Javier F, et al. Genome-wide assessment of Parkinson ’ s disease in a Southern Spanish population. Neurobiol Aging 2016; 45: 213.e3–213.e9.

7. Desikan RS, Schork AJ, Wang Y, et al. Genetic overlap between Alzheimer ’ s disease and Parkinson ’ s disease at the MAPT locus. Mol Psychiatry 2015; : 1588–95.

8. Nalls MA, Pankratz N, Lill CM, et al. Large-scale meta-analysis of genome-wide association data identifies six new risk loci for Parkinson’s disease. Nat Genet 2014; 46: 989–93.

9. Bandres-Ciga S, Ahmed S, Sabir MS, et al. The genetic architecture of Parkinson disease in Spain: characterizing population-specific risk, differential haplotype structures, and providing etiologic insight. Mov Disord 2019. DOI:10.1101/609016.

10. Nalls MA, Blauwendraat C, Vallerga CL, et al. Identification of novel risk loci , causal insights , and heritable risk for Parkinson ’ s disease : a meta-analysis of genome-wide association studies. Lancet Neurol 2019; : 1091–102.

11. Galvin JE, Pollack J, Morris JC. Clinical phenotype of Parkinson disease dementia. Neurology 2006; 67: 1605–12.

12. Aarsland D, Zaccai J, Brayne C. A Systematic Review of Prevalence Studies of Dementia in Parkinson ’ s Disease. Mov Disord 2005; 20: 1255–63.

13. Massano J, Bhatia KP. Clinical Approach to Parkinson’s Disease: Features, Diagnosis, and Principles of Management. Cold Spring Harb Perspect Med 2012; : 1–15.

14. Spillantini MG, Schmidt ML, Lee VM-Y, Trojanowski JQ, Jakes R, Goedert M. alpha-Synuclein in Lewy bodies. Nature 1997; 388: 839–40.

15. Dickson DW. Neuropathology of Parkinson Disease. Parkinsonism Relat Disord 2018; 46: 1–11.

16. Zhang X, Gao F, Wang D, et al. Tau Pathology in Parkinson ’ sDisease. Front Neurol 2018; 9: 1–7.

17. Espay AJ, Vizcarra JA, Marsili L, et al. Revisiting protein aggregation as pathogenic in sporadic Parkinson and Alzheimer diseases. Neurology 2019; 92: 329–37.

18. Zabetian CP, Hutter CM, Factor S a, et al. Association Analysis of MAPT H1 Haplotype and Subhaplotypes in Parkinson’s Disease. Ann Neurol 2007; 62: 137–44.

19. Vandrovcova J, Pittman AM, Malzer E, et al. Association of MAPT haplotype-tagging SNPs with sporadic Parkinson’s disease. Neurobiol Aging 2009; 30: 1477–82.

20. Boettger LM, Handsaker RE, Zody MC, McCarroll SA. Structural haplotypes and recent evolution of the human 17q21.31 region. 2012; 44: 881–5.

21. Steinberg KM, Antonacci F, Sudmant PH, et al. Structural diversity and African origin of the 17q21 . 31 inversion polymorphism. Nat Publ Gr 2012; 44: 872–80.

22. Brück D, Wenning GK, Stefanova N, Fellner L. Glia and alpha-synuclein in neurodegeneration: A complex interaction. Neurobiol Dis 2016; 85: 262–74.

23. di Domenico A, Carola G, Calatayud C, et al. Patient-Specific iPSC-Derived Astrocytes Contribute to Non-Cell-Autonomous Neurodegeneration in Parkinson’s Disease. Stem Cell Reports 2019; 12: 213–29.

24. Booth HDE, Hirst WD, Wade-Martins R. The Role of Astrocyte Dysfunction in Parkinson’s Disease Pathogenesis. Trends Neurosci 2017; 40: 358–70.

25. Sonninen TM, Hämäläinen RH, Koskuvi M, et al. Metabolic alterations in Parkinson’s disease astrocytes. Sci Rep 2020; 10. DOI:10.1038/s41598-020-71329-8.

26. Wakabayashi K, Hayashi S, Yoshimoto M, Kudo Yh, Takahashi H. NACP alphasynuclein-positive filamentous inclusions. Acta Neuropathol 2000; 99: 14–20.

27. Song YJC, Halliday GM, Holton JL, et al. Degeneration in different parkinsonian syndromes relates to astrocyte type and astrocyte protein expression. J Neuropathol Exp Neurol 2009; 68: 1073–83.

28. Braak H, Sastre M, Del Tredici K. Development of α synuclein immunoreactive astrocytes in the forebrain parallels stages of intraneuronal pathology in sporadic Parkinson’s disease. Acta Neuropathol 2007; 114: 231–41.

29. Lee HJ, Suk JE, Patrick C, et al. Direct transfer of α-synuclein from neuron to astroglia causes inflammatory responses in synucleinopathies. J Biol Chem 2010; 285: 9262–72.

30. Rothhammer V, Quintana FJ. Control of autoimmune CNS inflammation by astrocytes. Semin Immunopathol 2015; 37: 625–38.

31. Zhou Y, Zhu Y. Important role of the IL-32 inflammatory network in the host response against viral infection. Viruses 2015; 7: 3116–29.

32. Soutar MPM, Melandri D, Annuario E, et al. Regulation of mitophagy by the NSL complex underlies genetic risk for Parkinson’s disease at Chr16q11.2 and on the MAPT H1 allele. bioRxiv 2020. DOI:10.1101/2020.01.06.896241.

33. Giannuzzi G, Siswara P, Malig M, et al. Evolutionary dynamism of the primate LRRC37 gene family. Genome Res 2013; 23: 46–59.

34. Zody MC, Jiang Z, Fung H-C, et al. Evolutionary toggling of the MAPT 17q21.31 inversion region. Nat Genet 2008; 40: 1076–83.

35. Falola MI, Wiener HW, Wineinger NE, et al. Genomic Copy Number Variants: Evidence for Association with Antibody Response to Anthrax Vaccine Adsorbed. PLoS One 2013; 8. DOI:10.1371/journal.pone.0064813.

36. Miklossy J, Doudet DD, Schwab C, Yu S, McGeer EG, McGeer PL. Role of ICAM-1 in persisting inflammation in Parkinson disease and MPTP monkeys. Exp Neurol 2006; 197: 275–83.

37. Koprich JB, Reske-Nielsen C, Mithal P, Isacson O. Neuroinflammation mediated by IL-1β increases susceptibility of dopamine neurons to degeneration in an animal model of Parkinson’s disease. J Neuroinflammation 2008; 5: 1–12.

38. Zhang Y, Sloan SA, Clarke LE, et al. Purification and characterization of progenitor and mature human astrocytes reveals transcriptional and functional differences with mouse. Neuron 2016; 89: 37–53.

39. Huang A, Martin ER, Vance JM, Cai X. Detecting genetic interactions in pathway-based genome-wide association studies. Genet Epidemiol 2014; 38: 300–9.

40. Kong Y, Liang X, Liu L, et al. High throughput sequencing identifies MicroRNAs mediating α-synuclein toxicity by targeting neuroactive-ligand receptor interaction pathway in early stage of Drosophila Parkinson’s disease model. PLoS One 2015; 10: 1–24.

41. Fallon J, Reid S, Kinyamu R, et al. In vivo induction of massive proliferation, directed migration, and differentiation of neural cells in the adult mammalian brain. Proc Natl Acad Sci U S A 2000; 97: 14686– 91.

42. Mena MA, García De Yébenes J. Glial cells as players in parkinsonism: The ‘good,’ the ‘bad,’ and the ‘mysterious’ glia. Neuroscientist 2008; 14: 544–60.

43. Purcell S, Neale B, Todd-Brown K, et al. PLINK: A Tool Set for Whole-Genome Association and Population-Based Linkage Analyses. Am J Hum Genet 2007; 81: 559–75.

44. Consortium TIH. The International HapMap Project. Nature 2003; 426: 789–96.

45. Patterson N, Price AL, Reich D. Population structure and eigenanalysis. PLoS Genet 2006; 2: 2074–93.

46. Das S, Forer L, Schonherr S, et al. Next-generation genotype imputation service and methods Sayantan. 2016; 48: 1284–7.

47. Lumley T. rmeta: Meta-Analysis. R package version 3.0. 2018; : https://CRAN.R-project.org/package=rmeta.

48. Barrett JC, Fry B, Maller J, Daly MJ. Haploview: analysis and visualization of LD and haplotype maps. BIOINFORMATICS 2005; 21: 263–5.

49. Gabriel SB, DeFelice M, Rotimi C, et al. The structure of haplotype blocks in the human genome. Science (80- ) 2002; 296: 2225–9.

50. Aragon TJ. epitools: Epidemiology Tools. R package version 0.5-10. 2017; : https://CRAN.R-project.org/package=epitools.

51. Katz Y, Wang ET, Airoldi EM, Burge CB. Analysis and design of RNA sequencing experiments for identifying isoform regulation. Nat Methods 2010; 7: 1009–15.

52. Karch CM, Kao AW, Karydas A, et al. A Comprehensive Resource for Induced Pluripotent Stem Cells from Patients with Primary Tauopathies. Stem cell reports 2019; 13. DOI:10.1016/j.stemcr.2019.09.006.

53. Hoffman GE, Hartley BJ, Flaherty E, et al. Transcriptional signatures of schizophrenia in hiPSC-derived NPCs and neurons are concordant with post-mortem adult brains. Nat Commun 2017; 8. DOI:10.1038/s41467-017-02330-5.

54. Bowles KR, Julia TCW, Qian L, Jadow BM, Goate AM. Reduced variability of neural progenitor cells and improved purity of neuronal cultures using magnetic activated cell sorting. PLoS One 2019; 14: 1– 18.

55. Tcw J, Wang M, Pimenova AA, et al. An Efficient Platform for Astrocyte Differentiation from Human Induced Pluripotent Stem Cells. Stem cell reports 2017; 9: 600–14.

56. Patro R, Duggal G, Love MI, Irizarry RA, Kingsford C. Salmon: fast and bias-aware quantification of transcript expression using dual-phase inference. Nat Methods 2017; 14: 417.

57. Ritchie ME, Phipson B, Wu D, et al. limma powers differential expression analyses for RNA-sequencing and microarray studies. Nucleic Acids Res 2015; 43: e47.

58. Subramanian A, Subramanian A, Tamayo P, et al. Gene set enrichment analysis: a knowledge-based approach for interpreting genome-wide expression profiles. Proc Natl Acad Sci U S A 2005; 102: 15545– 50.

59. Reimand J, Kull M, Peterson H, Hansen J, Vilo J. G:Profiler-a web-based toolset for functional profiling of gene lists from large-scale experiments. Nucleic Acids Res 2007; 35: 1–8.

60. Raudvere U, Kolberg L, Kuzmin I, et al. g:Profiler: a web server for functional enrichment analysis and conversions of gene lists (2019 update). Nucleic Acids Res 2019; : 1–8.

61. Shannon P, Markiel A, Ozier O, et al. Cytoscape : A Software Environment for Integrated Models of Biomolecular Interaction Networks Cytoscape L Software Environment for Integrated Models of Biomolecular Interaction Networks. GENOM Les 2003; : 2498–504.

62. Merico D, Isserlin R, Stueker O, Emili A, Bader GD. Enrichment map: A network-based method for gene-set enrichment visualization and interpretation. PLoS One 2010; 5. DOI:10.1371/journal.pone.0013984.

